# Asymmetrical pegs in square holes? Functional and phylogenetic determinants of plant community assembly in temperate forest understories

**DOI:** 10.1101/2020.10.14.312512

**Authors:** Jared J. Beck, Daijiang Li, Sarah E. Johnson, David Rogers, Kenneth M. Cameron, Kenneth J. Sytsma, Thomas J. Givnish, Donald M. Waller

**Affiliations:** Department of Botany, University of Wisconsin-Madison, 430 Lincoln Dr, Madison, WI (USA); Department of Biological Sciences, Louisiana State University, Baton Rouge, LA (USA); Center for Computation & Technology, Louisiana State University, Baton Rouge, LA (USA); Department of Biology, Northland College, Ashland, WI (USA); Department of Biological Sciences, University of Wisconsin-Parkside, Kenosha, WI (USA)

**Keywords:** phylogeny, functional traits, coexistence, Wisconsin, environmental filtering, community assembly, resource partitioning

## Abstract

Despite advances in community assembly theory, uncertainties remain regarding how ecological and evolutionary processes shape species distributions and communities. We analyzed patterns of occurrence for 139 herbaceous plant species across 257 forest stands in Wisconsin (USA) to test predictions from community assembly theory. Specifically, we applied Bayesian phylogenetic linear mixed effects models (PGLMMs) to examine how functional traits and phylogenetic relationships influence plant distributions along environmental gradients and how functional similarity and phylogenetic relatedness affect local species co-occurrence. Leaf height, specific leaf area, and seed mass mediate species distributions along edaphic, climatic, and light gradients. In contrast, functional trait similarity and phylogenetic relationships only weakly affect patterns of local co-occurrence. These results confirm that broad-scale plant distributions are largely shaped by ecological sorting along environmental gradients but suggest deterministic assembly rules based on niche differentiation and complementary resource use may not govern local species co-occurrence in homogeneous environments.

**Statement of authorship:** JB conceived the idea for the study. DL, SJ, and DR collected the vegetation and functional trait data. JB analyzed the data with assistance from DL. KC, KS, TG, and DW secured funding for research and oversaw data collection. JB wrote the first draft of the manuscript, all authors contributed to manuscript revisions.

**Data accessibility statement:** Upon acceptance, data will be archived at Figshare (https://figshare.com/) and scripts used to analyze the data will be shared on Github (https://github.com/jaredjbeck/).

## INTRODUCTION

Disentangling the complex interplay of biotic and abiotic factors that structure ecological communities has been a central focus in ecology for more than a century (Clements 1916, Gleason 1926, Curtis 1959, MacArthur & Levins 1967, Tilman 1982, Weiher & Keddy 1999, Vellend 2016). Recent conceptual developments in coexistence theory coupled with methodological advances have stimulated renewed interest in community assembly (Cavender-Bares et al. 2009, Weiher & Freund 2011, HilleRisLambers et al. 2012). Much of this recent work examines how functional traits (Westoby & Wright 2006, McGill et al. 2006, Funk et al. 2017) and/or phylogenetic relationships (Webb et al. 2002, Vamosi et al. 2009, Cavender-Bares et al. 2009) affect community assembly. For example, we expect traits mediate species’ responses to environmental variation, leading organisms to occupy environments where they are best able to survive and compete for resources. This ecological sorting promotes co-occurrence among functionally similar species (Cornwell & Ackerly 2009, Kunstler et al. 2012, Kraft et al. 2015a). Trait similarities among species can reflect their shared evolutionary history and phylogenetic niche conservatism (Losos 2008, Wiens et al. 2010). Alternatively, trait similarities can also arise from evolutionary convergence on morphological or physiological characteristics reflecting selection for traits that confer competitive advantages under particular conditions (Cavender-Bares et al. 2006, Swenson et al. 2006, 2007). In either case, we expect strong trait-environment relationships lead to co-occurrence among functionally similar (and potentially related) species. Whereas ecological sorting promotes co-occurrence among functionally similar species, competition is expected favor coexistence among ecologically divergent species with complementary resource requirements (Tilman 1982, Chesson 2000, Mayfield & Levine 2010, Kraft et al. 2015b). Theory predicts limiting similarity leads to competitive exclusion among ecologically similar species. Such species would include close relatives if ecologically significant traits are phylogenetically conserved. Alternatively, closely related species may readily coexist when evolution has favored niche diversification within lineages (Givnish 1997)

Advancing our understanding of ecological communities hinges on reconciling these conflicting predictions about how trait differences influence community assembly and coexistence, including the spatial scales at which ecological convergence and niche differentiation operate (Cavender-Bares et al. 2009, Weiher & Freund 2011, HilleRisLambers et al. 2012). Many ecologists now favor a hierarchical model of community assembly in which ecological communities emerge from a set of nested processes operating at different spatial scales. This heuristic model asserts that local communities form as individual organisms from the regional species pool disperse and sort into communities based on their traits and ability to successfully compete within the local environmental context (Weiher & Keddy 1999, Kraft et al. 2015a). At finer scales within these communities, biotic interactions – especially competition and resource partitioning – are expected to favor local coexistence among species with different niches (Weiher & Keddy 1999, Silvertown et al. 2006, HilleRisLambers et al. 2012). Yet, unpredictable ecological interactions at local scales may preclude the evolution of such highly specialized resource niches (Janzen 1985), resulting in local communities shaped by competitive hierarchies (Mayfield & Levine 2010) or non-equilibrium processes driven by dispersal and ecological drift (Hubbell 2001; Alonso et al. 2006). So, while spatial scale-dependence may resolve the tension between ecological convergence and divergence in certain systems (Cavender-Bares et al. 2009, Scherrer et al. 2019), we still need to test the broader applicability of this hierarchical model of community assembly.

Here, we integrate extensive data on forest plant community composition, environmental conditions, functional traits, and a high-resolution molecular phylogeny to investigate how trait values and phylogenetic relationships affect species’ distributions along environmental gradients and within local communities across Wisconsin, U.S.A. We analyze data from a large set of temperate forest plant communities, focusing on the herbaceous taxa that comprise more than 80% of the plant species present in temperate forest plant communities (Gilliam 2007). Forest herbs share general characteristics well-adapted to forest understory conditions but also differ greatly in key morphological and physiological characteristics reflecting the diverse habitats they occupy (Bierzychudek 1982, Whigham 2004). For example, herb distributions respond sensitively to edaphic conditions and light availability (Curtis 1959, Givnish 1982, 1987, Gilliam 2014). Species turnover along these environmental gradients reflect physiological tradeoffs suggesting that environmental sorting and competitive hierarchies structure herb distributions at both local scales (Bratton 1976, Beatty 1984, Gilbert & Lechowicz 2004, Catella et al. 2019) and across landscapes (Curtis 1959, Amatangelo et al. 2014, Beatty 2014, Peet et al. 2014). The potential for trait differences to stabilize local coexistence among forest herbs has received less attention. Furthermore, we lack a comprehensive understanding of how phylogenetic relationships affect herb distributions and co-occurrence patterns. We therefore specifically ask: (1) To what extent do functional traits mediate plant responses to environmental variation? (2) Do shared responses to environmental variation lead to co-occurrence among closely related species (i.e. phylogenetic attraction)? (3) Are functionally similar or closely related species less likely to co-occur within forest stands or microsites because of limiting similarity or niche diversification (i.e. functional or phylogenetic repulsion)?

## MATERIALS AND METHODS

### Study region

We focus on 257 forest stands distributed across Wisconsin (and the western Upper Peninsula of Michigan – Fig. 1). Forest stands included in this study reflect four community types that differ in species composition, environmental conditions and ecological changes over the past 50 years (see citations listed). **Northern Upland Forests** (NUF) are dominated by sugar maple (*Acer saccharum*), eastern hemlock (*Tsuga canadensis*), and white pine (*Pinus strobus*) (Curtis 1959, Rooney et al. 2004). **Pine barrens** (PB) occur on sandy, well-drained soils dominated by xeric tree species like jack pine (*Pinus banksiana*) and northern pin oak (*Quercus ellipsoidalis*) (Habeck 1959, Li & Waller 2015). **Southern Upland Forests** (SUF) are dominated by oaks (*Quercus*) and hickories (*Carya*) in uplands and maples (*Acer*) or basswood (*Tilia americana*) on more fertile soils (Curtis 1959, Rogers et al. 2008). **Southern Lowland Forests** (SLF) are dominated by various species including silver maple (*Acer saccharinum*), ashes (*Fraxinus*), and elms (*Ulmus*) (Curtis 1959, Johnson & Waller 2013, Johnson et al. 2013, 2016).

**Figure 1.**
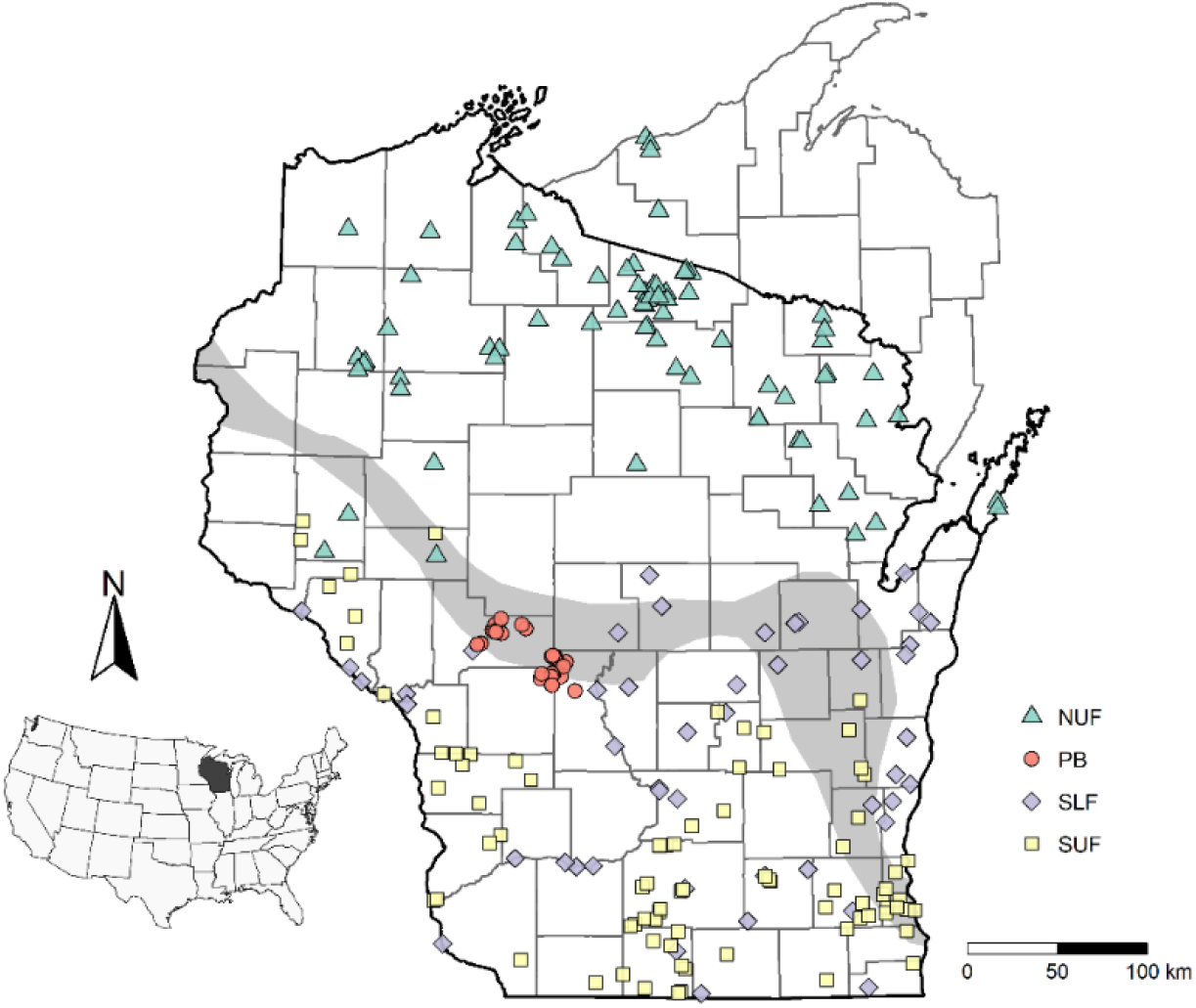
Map of study sites in Wisconsin, USA. Vegetation surveys were conducted at 257 forest stands across the state of Wisconsin between 2000 and 2012. Community classification follows Curtis (1959): blue triangles – Northern Upland Forests (NUF), red circles – Pine Barrens (PB), purple diamonds – Southern Lowland Forests (SLF), and yellow squares – Southern Upland Forests (SUF). The grey band illustrates the location of a pronounced floristic tension zone marking the range of limits of plant species with northern vs. southern affinities (see Fig. 2 and Curtis 1959).

### Vegetation sampling

To characterize herbaceous plant distributions and co-occurrence patterns, we analyzed extensive understory plant community survey data compiled in Wisconsin between 2000 and 2012 from stands originally surveyed between 1946 and 1958 (Curtis 1959). The resurveys were more intensive and obtained further data on overstory and soil conditions plus surrounding landscape composition and conditions (Rooney et al. 2004, Wiegmann & Waller 2006, Rogers et al. 2008, 2009, Waller et al. 2012, Johnson & Waller 2013, Johnson et al. 2013, 2016, Li & Waller 2015). Despite shifts in species composition over the 50+ year intervals, species-environment and trait-environment relationships have remained stable (Amatangelo et al. 2014). In resurveys, the presence of all vascular plant species was scored within 40-120+ replicate 1×1m quadrats (Table 1; see citations above for detailed field methods). We analyzed plant distributions and co-occurrence patterns for the 139 herbaceous plant species for which we have complete functional trait data (see below) and occurrence data in at least five stands (Fig. 2). These species represent 77% of all site-level occurrences and 84% of quadrat-level herb occurrences across these 257 forest sites.

**Figure 2.**
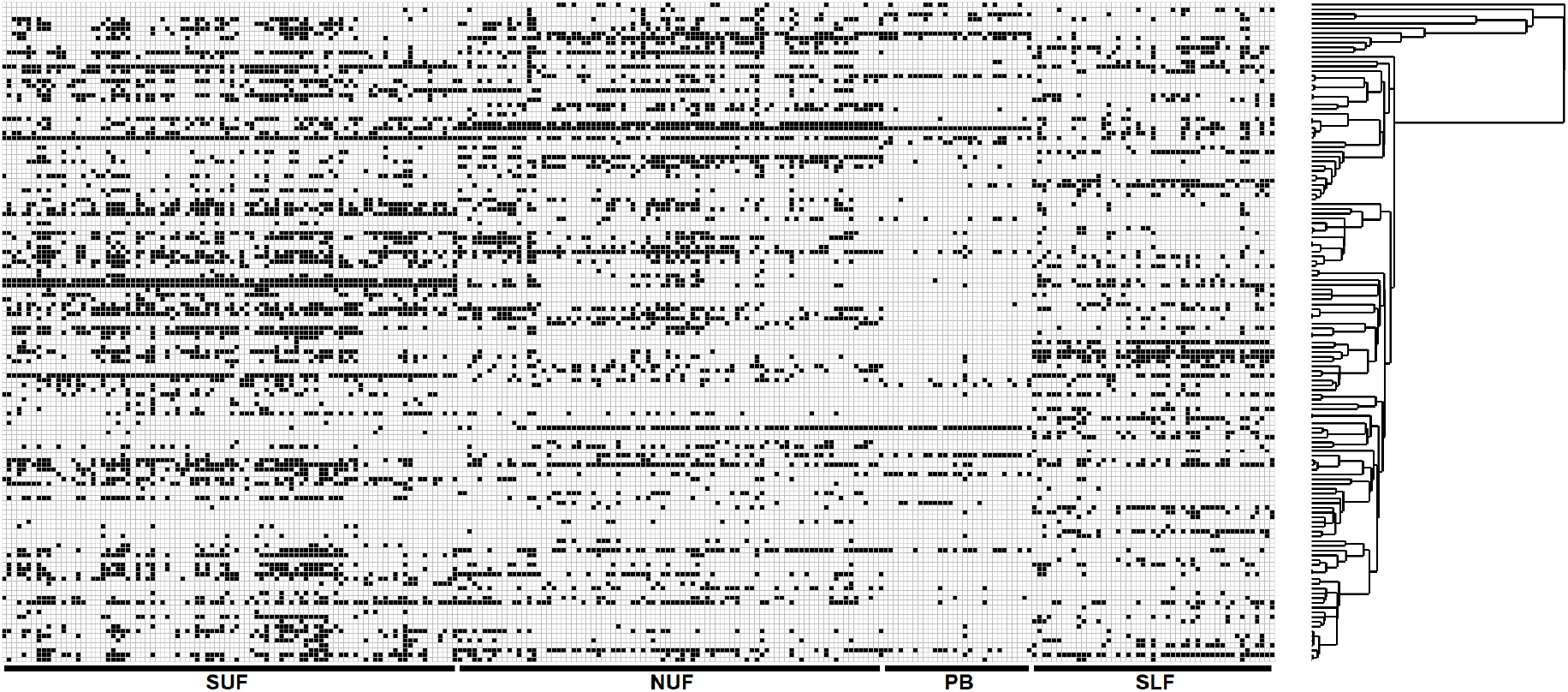
Heatmap illustrating distributional and co-occurrence pattern among 139 herbaceous species (rows) distributed across 257 study sites (columns) in Wisconsin (USA). Filled cells represent sites in which a particular species was present. The tree illustrates phylogenetic relationships among herbaceous plant species. Black bars and labels correspond to community types (Curtis 1959): SUF – Southern Upland Forest, NUF – Northern Upland Forest, PB – Pine Barrens, SLF – Southern Lowland Forest.

**Table 1.**
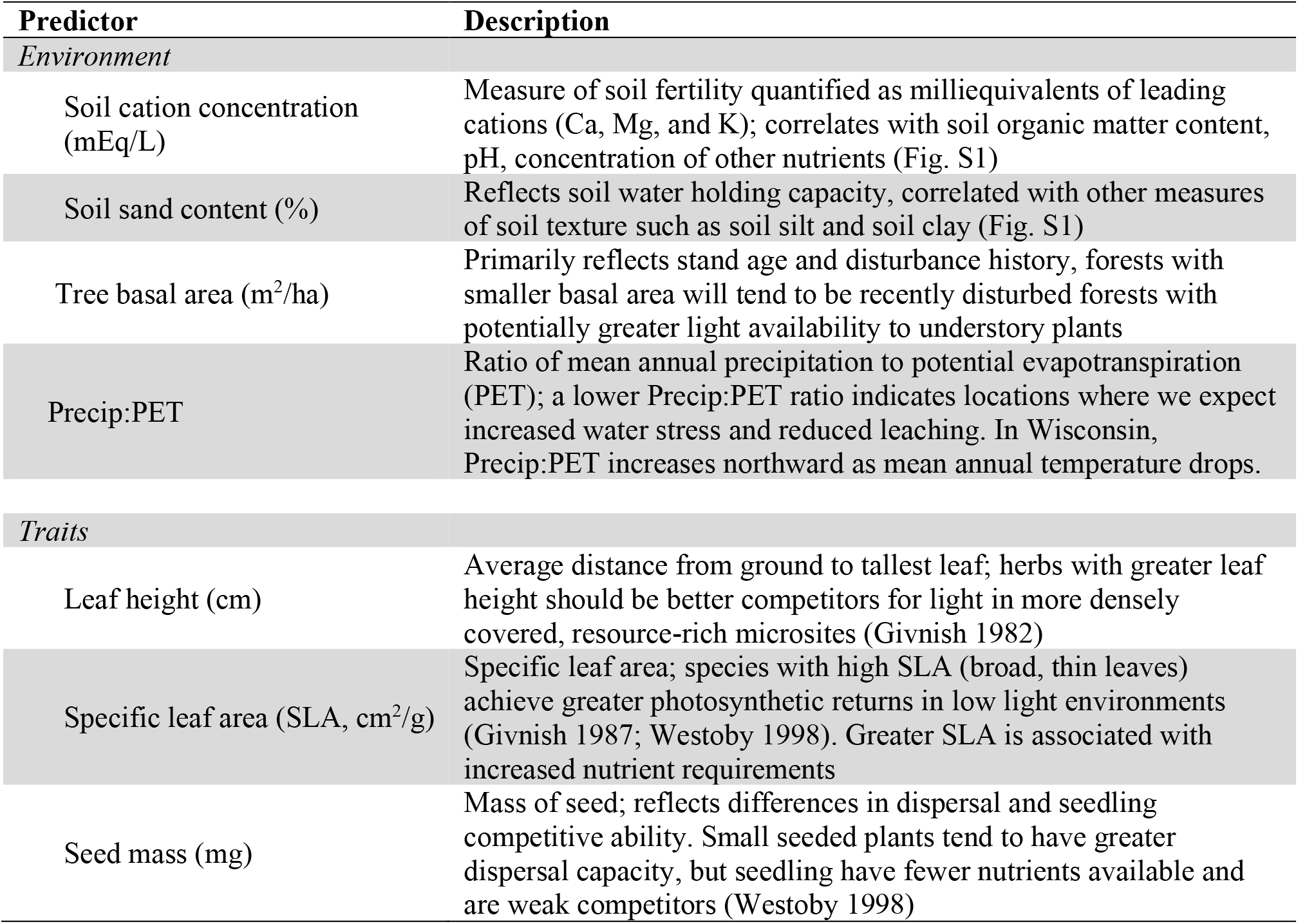
Description of environmental and trait predictors used in our phylogenetic generalized linear mixed effects models as well as expected influence on herb distributions.

| Predictor | Description |
| --- | --- |
| *Environment* |  |
| Soil cation concentration (mEq/L) | Measure of soil fertility quantified as milliequivalents of leading cations (Ca, Mg, and K); correlates with soil organic matter content, pH, concentration of other nutrients (Fig. S1) |
| Soil sand content (%) | Reflects soil water holding capacity, correlated with other measures of soil texture such as soil silt and soil clay (Fig. S1) |
| Tree basal area (m<sup>2</sup>/ha) | Primarily reflects stand age and disturbance history, forests with smaller basal area will tend to be recently disturbed forests with potentially greater light availability to understory plants |
| Precip:PET | Ratio of mean annual precipitation to potential evapotranspiration (PET); a lower Precip:PET ratio indicates locations where we expect increased water stress and reduced leaching. In Wisconsin, Precip:PET increases northward as mean annual temperature drops. |
| *Traits* |  |
| Leaf height (cm) | Average distance from ground to tallest leaf; herbs with greater leaf height should be better competitors for light in more densely covered, resource-rich microsites (Givnish 1982) |
| Specific leaf area (SLA, cm<sup>2</sup>/g) | Specific leaf area; species with high SLA (broad, thin leaves) achieve greater photosynthetic returns in low light environments (Givnish 1987; Westoby 1998). Greater SLA is associated with increased nutrient requirements |
| Seed mass (mg) | Mass of seed; reflects differences in dispersal and seedling competitive ability. Small seeded plants tend to have greater dispersal capacity, but seedling have fewer nutrients available and are weak competitors (Westoby 1998) |

### Environmental data

We selected a suite of functional traits and environmental predictors expected to influence herbaceous plant distributions (rationale in Table 1). These include four environmental predictors: milliequivalents of leading soil cations (Ca, Mg, and K), soil sand content (%), tree basal area (m^2^/ha), and the ratio of mean annual precipitation to potential evapotranspiration (Precip:PET). Soil cation concentrations and % soil sand represent two primary sources of edaphic variation: soil fertility and texture, respectively (Fig. S1 shows a PCA of all soil variables). In our analyses, we log-transformed milliequivalents of leading soil cations to improve the distribution of residuals.

Although we lacked direct measurements of light levels within these stands, we used tree basal area per hectare as a proxy for light availability. Our basal area estimates derive from sampling 40+ trees >10 cm in diameter (DBH – Waller et al. 2012). Young, recently disturbed forests typically have less basal area and higher understory light levels than more mature forests. Stands of a given age should also have lower basal area and greater light availability in drier climates and on less fertile soils.

We characterized climate variation using a measure of relative moisture supply (Fig. S2): mean annual precipitation divided by potential evapotranspiration (hereafter Precip:PET – http://www.cgiar-csi.org). We calculated Precip:PET from monthly 1-km scale WORLDCLIM historical climate data for the period 1970 to 2000 (Trabucco et al. 2008, Zomer et al. 2008, Fick & Hijmans 2017).

### Functional traits

We analyzed the effects of three functional traits known to affect plant distributions or local competitive ability: leaf height, specific leaf area (SLA), and seed mass (Givnish 1982, Westoby 1998, Westoby et al. 2002, Westoby & Wright 2006; see Table 1 for rationale). Because several non-seed plants (true ferns, lycophytes, and horsetails) lack true seeds, we assigned the minimum seed mass in the data set to these species reflecting the size of their functionally analogous diaspores. Estimates for 12 species derive from the Royal Botanic Gardens Key Seed Information Database (2020) but all others reflect seed mass measured on field-collected Wisconsin plants. We log-transformed seed mass and leaf height to improve residual distributions.

### Phylogeny

We used a molecular phylogeny assembled for use with the vascular flora of Wisconsin (Fig. S4) (Spalink et al. 2018, Givnish et al., *in press*). This phylogeny includes over 2300 native and introduced species, based on DNA sequences from seven chloroplast genes (*matK*, *rbcL*, *atpB*, *atpF-atpH*, *ndhF*, *rpl32, trnH-psbA*; see Spalink et al. 2018 for details). We pruned this regional phylogeny to obtain a tree for the 139 focal species analyzed in this study.

### Analyses

We formulated three sets of Bayesian phylogenetic generalized linear mixed-effects models (PGLMM) to examine how species distributions and co-occurrence patterns were influenced by environmental conditions, functional traits, and phylogenetic relationships (Ives and Helmus 2011, Li and Ives 2017). PGLMMs are a generalization of multilevel models designed to facilitate analyses of species’ distributions while accounting for phylogenetic covariance among species. The first and second set of models we fit (***PGLMM-1*** and ***PGLMM-2***) characterize plant distributions among forest sites while the third set (***PGLMM-3***) focuses on plant co-occurrences within sites. For *PGLMM-1* and *PGLMM-2*, we visually inspected species-environment and trait-environment relationships, checking for unimodal relationships. Species distributions relative to Precip:PET were all monotonic, but many species responded unimodally to soil fertility (milliequivalents of soil cations), soil texture (% sand), and tree basal area. Among traits, seed mass also responded unimodally to soil calcium and basal area. We therefore included quadratic terms for these relationships when analyzing species distributions.

The probability of species occurrence within sites was modeled using a binomial distribution. We analyzed presence/absence rather than abundance to facilitate model convergence. Our ***PGLMM-1*** model followed the form:

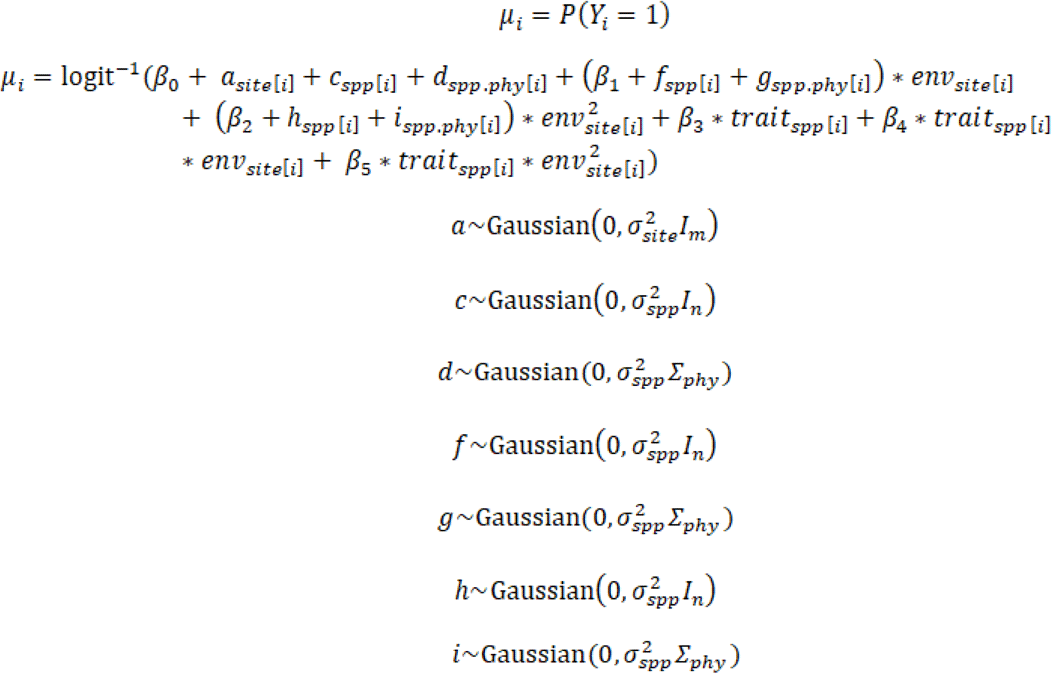

Where *β_0_* specifies the average probability of occurrence across all species, *β_1_* allows the probability of occurrence to vary according to an environmental predictor, *β_2_* accounts for non-monotonic species-environment responses, *β_3_* allows the average probability of plant occurrence to vary with the value of a trait, *β_4_* quantifies trait-environment interactions in which plant responses to environmental variation depend on trait differences, and *β_5_* accounts for unimodal trait-environment associations. *a* is a random effect that allows the overall probability of occurrence to vary among forest sites, *c* allows the probability of occurrence to vary among species (independent of phylogenetic relationships), *d* allows the probability of occurrence to vary among species such that there is phylogenetic covariance in species abundance, *f* allows species to respond differently to environmental variation (independent of phylogeny and traits), *g* allows for phylogenetic covariance in species responses to the environment, *h* allows for unimodal species responses to an environmental predictor, and *i* accounts for phylogenetic covariance in unimodal species-environment relationships.

We centered and scaled to unit variance all quantitative predictor variables to aid model convergence and to facilitate comparing predictors (Gelman & Hill 2007). For quadratic terms, we centered the predictor by subtracting its mean and then squared the centered estimates to minimize correlations between linear and quadratic predictors for the same environmental variable. Fixed effects were evaluated using 95% credible intervals. To assess random effects, we fit models with and without each random effect and compared the widely applicable information criterion (WAIC) of the two models. When the differences in WAIC (ΔWAIC) was greater than 5, we concluded that including the random effect substantially improved model performance (Spiegelhalter et al. 2002). In addition to the model including functional traits and trait-environment associations, we fit a second model using the same data and formulation but excluding traits and trait-environment associations. Comparing residual estimates these models allows us to quantify the extent to which functional traits drive phylogenetic patterns in species’ responses to environmental gradients (Li et al. 2017).

We applied ***PGLMM-2*** models to test for functional/phylogenetic attraction and repulsion among forest stands. Computational limitations limited our ability to model repulsion using the full dataset so we split this into two independent subsets (of 128 and 129 sites) and fit separate models (***PGLMM-2.1*** and ***PGLMM-2.2***). These subsets reflect a random sample of sites stratified by community. ***PGLMM-2*** models have the same structure and use the same environmental and functional trait predictors as *PGLMM-1* but retain random effects representing variation in occurrence probabilities across sites (term *a*), across species (*c*), and across species where occurrence probability is similar among close relatives (*d*). With these ***PGLMM-2*** models, we further compared reduced models to full models that included an additional covariance matrix specifying either functional repulsion, phylogenetic repulsion, functional attraction, and phylogenetic attraction (Ives and Helmus 2011). To obtain the phylogenetic attraction matrix, we scaled the phylogenetic covariance matrix so that its determinant equaled 1. We obtained the phylogenetic repulsion matrix by calculating the inverse of this attraction matrix. For both attraction and repulsion matrices, we then calculated the Kronecker product of species covariance matrices and a diagonal site matrix to obtain the covariance matrix blocked by site specifying phylogenetic attraction or repulsion among species that co-occur in forest stands. The functional attraction and repulsion matrices used functional traits to generate a dissimilarity matrix based on Gower distances. We calculated this dissimilarity matrix using a broader set of traits that could influence community dynamics including leaf height, SLA, leaf C:N, seed mass, longevity (short-lived vs. long-lived), pollination syndrome (biotic vs. abiotic), and flowering season (spring, summer, or fall). We then used a hierarchical clustering algorithm to produce a trait dendrogram and functional covariance matrix (Petchey & Gaston 2006; Fig. S5). As with phylogenetic covariance matrices, we scaled the functional covariance matrix so that its determinant equaled 1, found the inverse to obtain a functional repulsion matrix, and then calculated the Kronecker product to obtain the functional attraction and repulsion matrices blocked by site for use in our model. To assess the strength of functional/phylogenetic attraction and repulsion, we compared WAIC values from PGLMM models that included these covariance matrices to the reduced model excluding covariance matrices. As with *PGLMM-1*, we fit variations of *PGLMM-2* models that either included or excluded trait predictors and trait-environment interactions to evaluate whether traits and phylogenetic relationships provide complementary information about herb distributions.

Our final set of ***PGLMM-3*** models tested the strength of functional and phylogenetic associations at finer spatial scales by examining species co-occurrence patterns among the individual 1m^2^ quadrats within stands. These models were formulated as described above but included only trait predictors. For each site, we fit a reduced model that included a suite of trait predictors (leaf height, SLA, and seed mass) and random effects allowing variation in the probability of occurrence among quadrats, the probability of occurrence among species, and the probability of species occurrence conditioned on phylogenetic relatedness (analogous to the random effects *a*, *c*, and *d* described above). We then compared this reduced model to full models that added covariance matrices specifying either functional repulsion, phylogenetic repulsion, functional attraction, or phylogenetic attraction at the scale of quadrats. Comparing WAIC values from the full and reduced models allows us to evaluate the strength of functional and phylogenetic association for herb assemblages within forest stands. As in *PGLMM-1* and *PGLMM-2*, we also fit variations of *PGLMM-3* models that excluded trait predictors. All analyses were performed in R (R Core Team 2016) using ‘ggplot2’ (Wickham 2009) to generate figures. We fit PGLMMs using INLA (Rue et al. 2009) via the *communityPGLMM()* function within the ‘phyr’ package (Li et al., *in press*).

## RESULTS

Herbaceous plant distributions reflect variation in environmental conditions, functional traits, and often their interactions (Fig. 3). Overall, the probability of herb occurrence at sites with higher Precip:PET paralleling a general south-to-north decline in herb abundance and richness. Occurrence probability also increased with more leaf area per unit biomass (SLA). Overall occurrence probabilities first increased, then declined, as soils became more fertile (milliequivalents of leading cations) or sandier and as tree basal area increased. Probabilities of species occurrence tended to be higher for species with larger diaspores and lower for species with greater leaf height (though credible intervals just overlapped zero).

**Figure 3.**
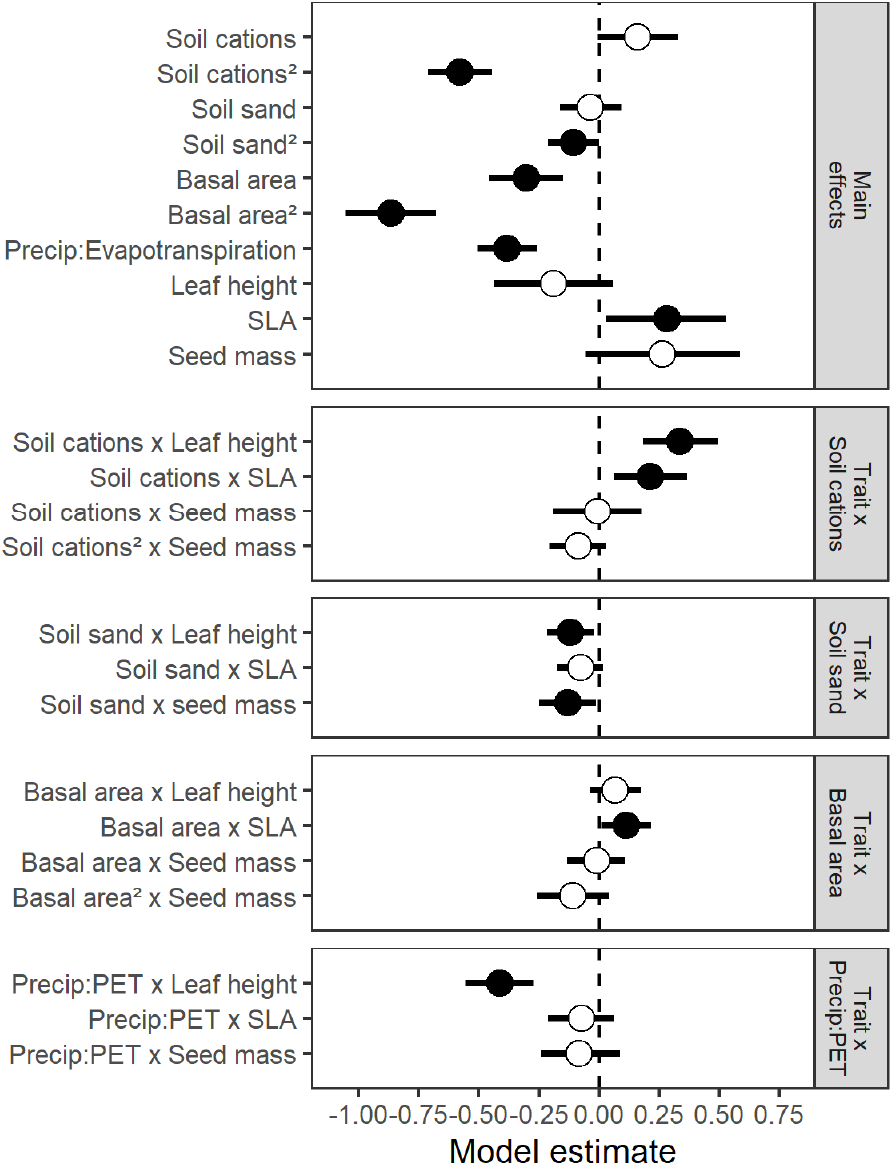
Estimated PGLMM coefficients for the effect of trait and environmental predictors on the probability of plant species occurrence. Top panel depicts main effects of predictors while the bottom four panels illustrate estimates for trait-environment interactions. Positive estimates indicate a positive relationship between predictors and the probability of occurrence. Solid circles indicate estimates that are different from zero (95% credible interval does not overlap with zero).

Herbaceous plant distributions along environmental gradients often reflect variation in key functional traits (Fig. 3). As soil fertility increased, the probability of encountering taller species increased. More fertile sites also supported species with greater SLA. Seed mass did not appear to respond to soil fertility. Sites with sandier soils favored shorter statured species with smaller seeds. SLA does not appear to mediate plant responses to soil texture. As expected, shadier understories (with higher basal area) support species with greater SLA. Leaf height and seed mass did not affect herb responses to variation in basal area among forest stands. Sites with greater Precip:PET support shorter plants regardless of their SLA or seed mass.

After accounting for the effects of traits and trait-environment relationships, we examined how residual variation can be attributed to sites, species, and phylogenetic relationships. Overall occurrence probabilities varied considerably among species. Herb distributions along gradients in soil fertility, soil sand, tree basal area, and Precip:PET also varied among species in ways that did not depend on the functional traits included in our model (Table S2; Figs. S5-S8). Many species responded unimodally to soil fertility, soil texture, and basal area, reaching their peak probability of occurrence at different points along environmental gradients (Figs. S5-S7). Including traits and trait-environment interactions generally reduced the amount of residual variation in species’ responses to environmental gradients (Fig. 4; Table S2). However, PGLMMs including traits provided comparable estimates to those excluding traits (Pearson correlation of deviation estimates: r = 0.99, N = 17, P < 0.001) and there was little evidence that closely related herbs responded similarly to environmental gradients whether PGLMMs included traits and trait-environment interactions or not (Table S2).

**Figure 4.**
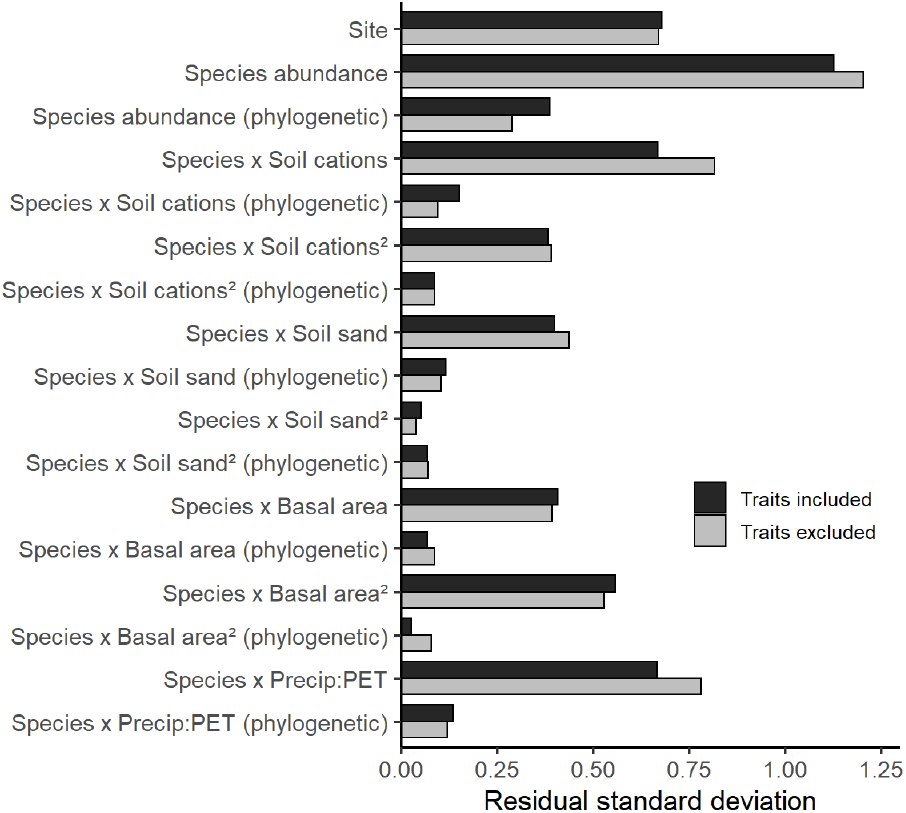
Estimated standard deviation for random effects in PGLMMs that include traits (black bars) and PGLMMs that exclude traits (grey bars). These estimates quantify residual variation in the probability of occurrence among sites, species, and differences in species’ distributions along environmental gradients. Random effects for species prevalence and species-environment association include terms quantifying variation independent of phylogenetic relationships and the degree to which phylogenetically related species exhibit similar abundance or responses to environmental variation (phylogenetic terms). We fit PGLMMs characterizing plant distributions across forest stands including traits and trait-environment interactions (PGLMM-1a – dark grey bars) as well as models excluding traits (PGLMM-1b – light grey bars) to quantify the extent to which functional traits drive phylogenetic patterns (Li and Ives 2017). Estimates of residual standard deviation from PGLMM-1a and PGLMM-1b were highly correlated (r = 0.99, N = 17, P < 0.001). See Table S2 for ΔWAIC values for each random effect term and summary of how including traits affected inferences.

Limiting similarity should discourage coexistence among ecologically similar species, but we found little evidence for functional repulsion in species co-occurrence patterns in our models, either among sites (Table S3) or within sites (Fig. 5; Fig. S9; Table S4). Site-level analyses (PGLMM-2) showed no indication that functionally similar species segregated among sites (ΔWAIC values for independent subsets: −1.34 and −1.51). There was some evidence that closely related species segregated among forest stands when traits and trait-environment interactions were included (ΔWAIC = 3.43 and 5.16), but these patterns were relatively weak. In contrast, we found strong evidence that functionally similar (ΔWAIC = 262.54 and 299.98) and phylogenetically related species co-occurred within forest stands (ΔWAIC = 61.39 and 160.35). At finer spatial scales within forest stands, we found weak evidence of local functional and phylogenetic repulsion (PGLMM-3). Functionally similar species showed clear evidence (ΔWAIC > 5) of quadrat-scale segregation within only 14 of 257 sites (5.4% - Fig. 5a). Closely related species segregated among microsites in just 14 sites (Fig. 5b). Meanwhile, we found evidence of functional attraction at 54 sites (21.0% - Fig. 5c) and phylogenetic attraction at 55 sites (21.4% - Fig. 5d). Including trait predictors generally increased the strength of residual functional and phylogenetic patterns (especially phylogenetic repulsion), but qualitative inferences generally remained similar regardless of whether traits were included as predictors in PGLMM-2 and PGLMM-3 (Tables S3, S4; Figs. S10, S11).

**Figure 5.**
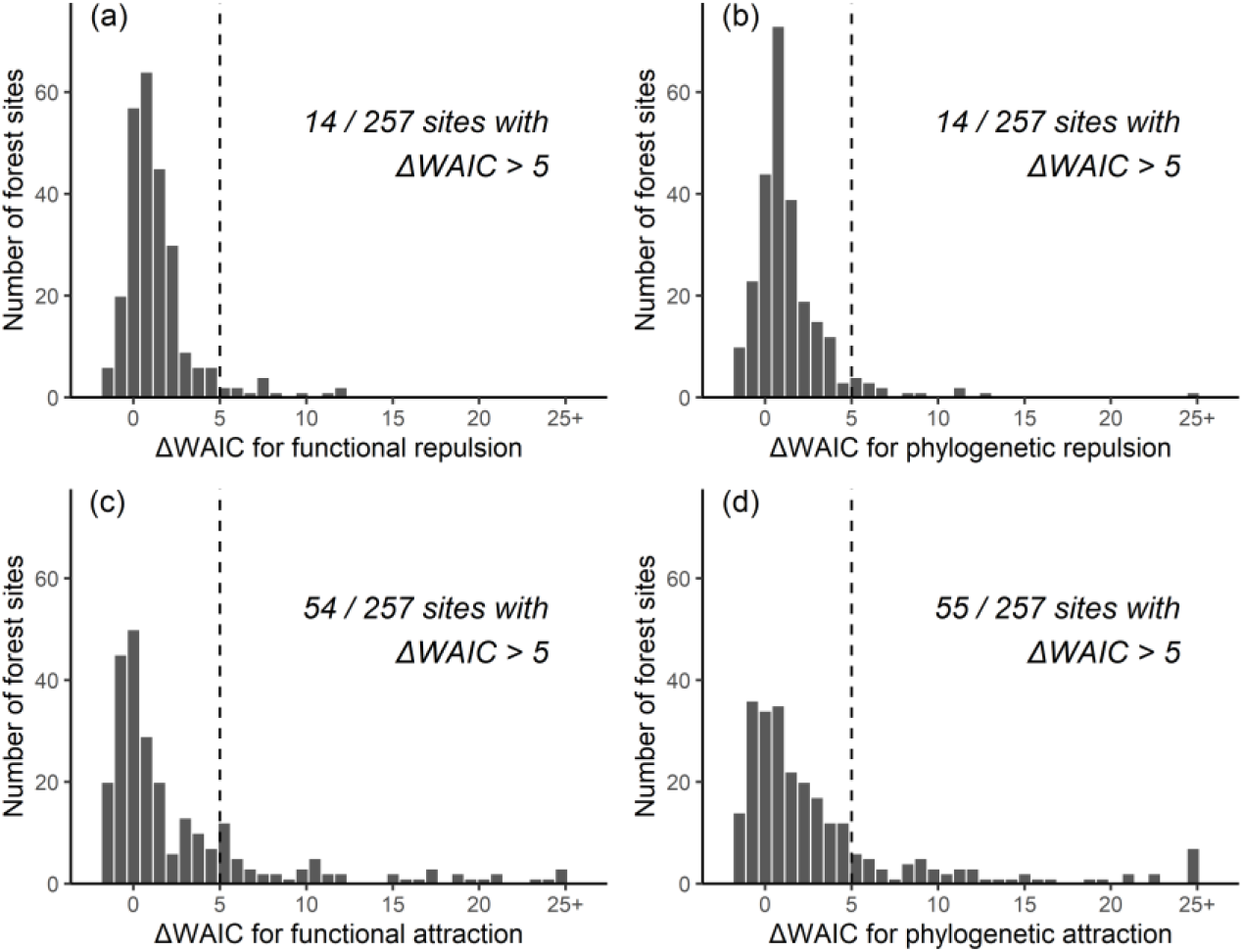
Patterns of (a) functional repulsion, (b) phylogenetic repulsion, (c) functional attraction, and (d) phylogenetic attraction at local scales within forest stands. These histograms display ΔWAIC values (the differences in WAIC values between PGLMMs with and without covariance matrices specifying repulsion or attraction, see PGLMM-3 in Methods) for all 257 forest sites. Only ΔWAIC values >5 (vertical dashed line) provide strong evidence of functional/phylogenetic repulsion (spatial segregation of functionally similar or phylogenetically related species among microsites within forest stands) or functional/phylogenetic attraction (spatial aggregation within microsites). Negative ΔWAIC values reflect cases where the penalty for additional complexity is greater than the amount of additional information gained by accounting for functional or phylogenetic repulsion.

## DISCUSSION

Here, we tested key predictions from community assembly theory using extensive data on the distribution and functional characteristics of 139 herbaceous plant species distributed over 257 temperate forest sites spanning 160,000+ km^2^. We used environmental data from all sites to explore ecological processes affecting the distribution of these species at multiple spatial scales. Consistent with previous studies from temperate forests and other ecosystems, we found that traits often mediate how plant species respond to landscape-scale environmental gradients in soil fertility, soil texture, light, and climate (Ordoñez et al. 2009, Pollock et al. 2012, Jamil et al. 2013, Amatangelo et al. 2014). Many of these responses are consistent with economic theory and previous studies of plant ecophysiology. For example, we found taller plants at sites with higher soil fertility (Fig. 3), as expected given that optimal leaf height increases in sites where higher soil fertility supports denser plant coverage (Givnish 1982, 1995). Greater soil fertility and light in southern lowland forests are associated with high densities of taller herbaceous plants (Menges & Waller 1983). Greater soil fertility should also increase optimal photosynthetic capacity, favoring plants with greater leaf N and SLA (Givnish 1979, Mooney & Gulmon 1982); increases in photosynthetic capacity should also increase height (Givnish 1995). In contrast, dry, nutrient-poor soils (such as those found in sandy pine barrens sites) limit plant growth, reducing herb coverage and favoring shorter species. Shadier (higher basal area) forests favored species with broader, thinner leaves (higher SLA - Fig. 3), reflecting the advantage of broader, thinner leaves in low light environments (Givnish 1979). Overall, soils in northern upland forests were sandier and less fertile than soils in southern forests (Fig. S1). Well-drained, infertile soils in northern upland forests (Fig. S1) reflect greater leaching (high Precip:PET), less soil development on glacially-derived sediments (i.e., glacial till and sandy outwash), and infertile bedrock (mostly sandstones, granites, and schists) compared to the calcareous bedrock prevalent in southern Wisconsin (Fig. S2; Mudrey et al. 1982). Small-seeded plants were more abundant at sites with sandy soils likely reflecting the fact that disturbed, sparsely vegetated environments with little competition favor small-seeded plants (Westoby 1998, Westoby & Wright 2006). These strong trait-environment associations re-affirm the importance of ecological sorting and competitive hierarchies in shaping local plant communities and plant distributions along environmental gradients (Curtis 1959; Whittaker 1956; Kraft et al. 2015a).

Although functional traits strongly influenced plant responses to environmental variation, plant distributions along environmental gradients were unrelated to phylogenetic relationships. Including trait-environment predictors in PGLMMs had little effect on phylogenetic patterns along environmental gradients (Fig. 4; Table S2), despite the strong phylogenetic signal in leaf height, SLA, and seed mass (Table S5). Complex patterns of trait evolution among these species likely cause the weak association between environmental conditions and phylogenetic relationships. Distantly related clades within the same environment often exhibit similar leaf shape, physiognomy, and phenology reflecting convergent evolution for traits that confer competitive advantages in particular environments (Givnish 1987, 1988, 1995). Conversely, divergent evolution and specialization for distinct habitats likely drive the substantial trait variation within clades such as Asteraceae and Poaceae as well as genera like *Viola* (Givnish 1987). Thus, the link between functional traits and phylogenetic relatedness in forest herbs is complicated by patterns of phylogenetic conservatism conflated with ecological convergence among distantly related clades and niche diversification within certain lineages. Despite some evidence that phylogenetically related species segregate among forest stands (perhaps reflecting niche diversification among close relatives), phylogenetic attraction was stronger (Table S3). This suggests unmeasured environmental variables, biogeographic history, or other biotic processes that promote co-occurrence among close relatives likely influence herb distributions.

Phylogenetic relatedness is often used as a surrogate for functional similarity under the assumption that ecologically relevant traits are conserved within evolutionary lineages (see Introduction). While evolutionary history clearly influences community assembly in these forest herbs, complex patterns of trait evolution and varied biogeographic histories confound efforts to devise simple heuristic approaches like equating phylogenetic relatedness to functional similarity. Relying on phylogenetic relationships to represent the morphological and physiological traits that mediate plant responses to environmental variation may work better at broad spatial scales where phylogenetic signals better reflect biogeographic history (Swenson & Enquist 2009). Such assumptions may also be valid within species-rich clades where stronger ecological interactions drive niche diversification (Cavender-Bares et al. 2006, 2018, Swenson et al. 2006, 2007, Helmus et al. 2007, Cavender-Bares 2019). Phylogenetic relationships appear less useful for explaining community assembly of herbaceous species at local or regional scales involving taxa from diverse clades like this study.

Contemporary community assembly theory predicts that after species sort along environmental gradients, biotic interactions (especially those associated with conflicting or complementary resource use) shape local community dynamics (Weiher & Keddy 1999, Cavender-Bares et al. 2009, HilleRisLambers et al. 2012). We expect limiting similarity to cause competitive exclusion among functionally similar and/or phylogenetically related species. However, we found little evidence for segregation among functionally similar species among communities or within sites, the scale where these herbs directly compete for resources. Evidence for phylogenetic repulsion was weak and emerged only among forest stands, likely reflecting diversification into different habitats. What accounts for the lack of niche differentiation among locally co-occurring herbs? The prevalence of rare species may reduce statistical power to detect functional or phylogenetic repulsion, especially when relying on presence-absence data.

Alternatively, phylogenetic relationships and the traits we analyzed may not have captured complementary ecological strategies stabilizing local coexistence within microsites (e.g. differences in rooting depth). Static co-occurrence patterns could also obscure relevant community dynamics. There is, however, little evidence for complementary resource usage among temperate forest herbs in the absence of spatial or temporal environmental heterogeneity (Amarasekare 2003, Silvertown 2004, Costanza et al. 2011, Beatty 2014, Catella et al. 2019). In contrast, differential plant responses to spatial environmental heterogeneity strongly influence local plant distributions (Beatty 2014). Fine-scale variation in microtopography, light levels, and soil depth, fertility, and moisture can all affect local patterns of distribution and co-occurrence (Bratton 1976, Thompson 1980, Beatty 1984, Crozier & Boerner 1984, Beck & Givnish, *in press*). In this study, the tendency for functionally similar and phylogenetically related herbs to occupy similar microsites could reflect local environmental heterogeneity and ecological sorting within sites (Table S4).

The fact that we only found evidence for functional and phylogenetic attraction in <30% of forest stands suggests that deterministic assembly rules may not exert a strong influence on local plant distributions. Previous studies of fine-scale co-occurrence patterns in these forests that ignored traits and phylogenetic relationships revealed weak interspecific associations (Li & Waller 2016). Recent studies investigating the potential for scale-dependent processes to shape plant community assembly have mostly focused on complementary resource use at scales where plants directly compete for resources (Maire et al. 2012, Gross et al. 2013, Scherrer et al. 2019). We hypothesize instead that strong ecological sorting across sites increases the role of equalizing processes within communities with relatively homogeneous environmental conditions (Chase 2014). Whereas stabilizing processes promote coexistence by reducing interspecific competition, equalizing processes reduce absolute fitness differences among species (Chesson 2000, Mayfield & Levine 2010). Thus, ecological sorting generates communities composed of species similarly well-equipped to compete for resources. This reduces competitive differences among co-occurring species which slows the rate of competitive exclusion while increasing the importance of local dispersal and stochastic demographic events. Although locally occurring species in our study differ ecologically (violating the assumption of ecological equivalence), reduced fitness differences among co-occurring species may prevent local communities from reaching equilibrium and cause community dynamics to resemble neutral communities driven by ecological drift (Hubbell 2001, Alonso 2006, Chase 2014). Equalizing processes may also set the stage for other relatively weak processes to influence local community dynamics such as differential responses to temporal environmental variation (Adler et al. 2006, Angert et al. 2009), demographic differences and life-history tradeoffs (Grubb 1977, Shmida & Ellner 1984), and localized plant-soil and plant-pathogen feedbacks (Bever et al. 2015, Smith & Reynolds 2015). Closer study of these stabilizing processes could yield further insights into fine-scale drivers of community assembly.

We conclude that trait-mediated ecological sorting and competitive hierarchies shape plant distributions along landscape-scale environmental gradients in many temperate forests. This ecological sorting results in local co-occurrence among functionally similar species from a variety of evolutionary lineages. These findings are largely consistent with predictions from community assembly theory and echo century-old concepts that emphasize how plant species sort individualistically into communities according to their relative competitive ability in a local environment (Gleason 1926, Whittaker 1956, Curtis 1959). The absence of strong functional or phylogenetic repulsion among co-occurring species in our study, however, departs from common predictions made by modern community assembly theory about the importance of niche differentiation and complementary resource use for local species coexistence. We hypothesize this strong environmental sorting which favors local co-occurrence among functionally similar species increases the importance of equalizing processes within relatively homogenous environments. While complementary resource use is often implicated in local species coexistence, we surmise that weak and/or unpredictable species interactions caused by the prevalence of rare species, divergent biogeographic histories, individualistic responses to environmental conditions, and local dispersal may preclude the evolution of complementary resource niches (the α-niche *sensu* Silvertown et al. 2006). Absent highly specialized resource niches, species may occupy microsites and niche space within local communities opportunistically, fitting themselves into local communities as “asymmetrical pegs in square holes” (Janzen 1985). Rather than looking exclusively to deterministic assembly rules based on niche differentiation and complementary resource use, we encourage ecologists to consider a broader suite of ecological processes that may shape the local distribution, abundance, co-occurrence, and diversity of plant species. These include specialized interactions with herbivores, pathogens, and other microbes as well as life-history tradeoffs associated with demographic differences in fecundity, longevity, and dispersal. Evaluating these processes and how their influence varies with spatial scale should yield new insights and a more synthetic understanding of plant community assembly.

## ACKNOWLEDGEMENTS

A. Ives, M. Turner, and B. Larget provided feedback that greatly improved the manuscript. A. Ives and R. Dinnage offered valuable statistical advice and assistance. The vegetation surveys and studies of trait variation were supported by a National Research Initiative Grant (2008-35320-18680) from the USDA National Institute of Food and Agriculture Biology of Weedy and Invasive Species Program as well as National Science Foundation grants DEB-023633, DEB-0717315, and a Dimensions of Biodiversity award (DEB-1046355) that also supported the phylogeny work. J. Beck was supported as an NSF Graduate Research Fellow (DGE-1747503). Opinions, findings, and conclusions expressed here are those of the authors and do not reflect the views of the NSF. Support was also provided by the Graduate School and the Office of the Vice Chancellor for Research and Graduate Education at the University of Wisconsin-Madison with funding from the Wisconsin Alumni Research Foundation.

---

**Table S1.** Summary of vegetation data sets used in analysis of community composition. Year of the vegetation surveys, the number of sites surveyed (range if applicable), the number of 1 x 1 meter quadrats sampled per site, and the number of taxa included in our analysis (139 unique taxa)

| Community type | Abbreviation | Survey year | No. sites | No. quadrats/site | No. taxa |
| --- | --- | --- | --- | --- | --- |
| Pine Barrens | PB | 2012 | 30 | 50 | 49 |
| Northern upland forest | NUF | 2000-04 | 86 | 80-120 (40-120) | 109 |
| Southern lowland forest | SLF | 2007-08 | 49 | 42 (39-42) | 109 |
| Southern upland forest | SUF | 2002-03 | 92 | 80 (40-403) | 115 |

**Table S2.** Random effects summary for PGLMM examining plant distributions across 257 forest sites (PGLMM-1 in Methods; Fig. 4). The table includes estimated standard deviation estimates and the difference in WAIC (ΔWAIC) between PGLMMs with and without each random effect term. ΔWAIC values for phylogenetic effects were estimated by comparing WAIC between the full model and a reduced model excluding phylogenetic covariance matrix for the term of interest. Non-phylogenetic random effects were evaluated by comparing the aforementioned model (excluding phylogenetic covariance) to a reduced model excluding the non-phylogenetic term of interest. A ΔWAIC greater than 5 indicates that including the random effect substantially improved PGLMMs. ΔWAIC values less than 0 indicates the penalty assessed when fitting a more complex model exceeds the information gained.

| Model term | Biological interpretation | Including traits |  | Excluding traits |  |
| --- | --- | --- | --- | --- | --- |
| | | $\sigma$ | ΔWAIC | $\sigma$ | ΔWAIC |
| Species abundance | Variation in occurrence probability among species | 1.126 | 2818.27 | 1.205 | 2891.48 |
| Species abundance (phylogenetic) | Related species have similar occurrence probabilities | 0.389 | -1.83 | 0.290 | 0.07 |
| Species x Soil cations | Herb species distributions respond differently to soil fertility | 0.670 | 326.52 | 0.817 | 402.09 |
| Species x Soil cations (phylogenetic) | Distribution of related species respond similarly to soil fertility | 0.153 | -5.13 | 0.097 | -2.29 |
| Species x Soil cations<sup>2</sup> | Variation in the strength of unimodal soil fertility distributions among herbs | 0.385 | 98.96 | 0.393 | 98.19 |
| Species x Soil cations<sup>2</sup> (phylogenetic) | Related species exhibit similar unimodal soil fertility distributions | 0.088 | -3.40 | 0.087 | -0.24 |
| Species x Soil sand | Herb species distributions respond differently to soil texture | 0.400 | 280.73 | 0.438 | 311.31 |
| Species x Soil sand (phylogenetic) | Distribution of related species respond similarly to soil texture | 0.117 | -3.15 | 0.104 | -0.86 |
| Species x Soil sand<sup>2</sup> | Variation in the strength of unimodal soil texture distributions among herbs | 0.052 | 16.92 | 0.039 | 16.35 |
| Species x Soil sand<sup>2</sup> (phylogenetic) | Related species exhibit similar unimodal soil texture distributions | 0.069 | 2.76 | 0.070 | 5.51 |
| Species x Basal area | Herb species distributions respond differently to basal area | 0.408 | 95.65 | 0.393 | 99.02 |
| Species x Basal area (phylogenetic) | Distribution of related species respond similarly to basal area | 0.068 | -5.04 | 0.088 | -1.12 |
| Species x Basal area<sup>2</sup> | Variation in the strength of unimodal basal area distributions among herbs | 0.559 | 79.60 | 0.529 | 80.41 |
| Species x Basal area<sup>2</sup> (phylogenetic) | Related species exhibit similar unimodal basal area distributions | 0.028 | -4.94 | 0.080 | -1.72 |
| Species x Precip:PET | Herb species distributions respond differently to Precip:PET | 0.667 | 699.21 | 0.782 | 949.23 |
| Species x Precip:PET (phylogenetic) | Distribution of related species respond similarly to Precip:PET | 0.136 | -2.94 | 0.121 | -0.281 |

**Table S3.** Summary information for site-level models quantifying the strength of residual functional and phylogenetic co-occurrence patterns (see **PGLMM-2** in Methods). We fit PGLMMs for two independent sets of forest sites (Subset 1 and Subset 2 respectively) and compared models with and without covariance matrices specifying site-level functional repulsion, phylogenetic repulsion, functional attraction, and phylogenetic attraction. The table includes standard deviation estimates quantifying the amount of residual variation attributed to functional and phylogenetic repulsion or attraction as well as the difference in WAIC (ΔWAIC) between PGLMMs with and without each repulsion covariance matrix. A ΔWAIC greater than 5 indicates that including the covariance matrix substantially improved models of species distributions and co-occurrence patterns. ΔWAIC values less than 0 indicates the penalty assessed when fitting a more complex model exceeds the information gained.

| Model term | Biological interpretation | Including traits |  | Excluding traits |  |
| --- | --- | --- | --- | --- | --- |
| | | $\sigma$ | ΔWAIC | $\sigma$ | ΔWAIC |
| Subset 1 (PGLMM-2.1) |  |  |  |  |  |
| Functional repulsion | Functionally similar species segregate among forest stands | 0.012 | -1.34 | 0.011 | -1.89 |
| Phylogenetic repulsion | Phylogenetically related species segregate among forest stands | 0.025 | 3.43 | 0.011 | 0.97 |
| Functional attraction | Functionally similar species aggregate within forest stands | 0.285 | 262.54 | 0.349 | 564.17 |
| Phylogenetic attraction | Phylogenetically related species aggregate within forest stands | 0.172 | 61.39 | 0.145 | 51.00 |
| Subset 2 (PGLMM-2.2) |  |  |  |  |  |
| Functional repulsion | Functionally similar species segregate among forest stands | 0.013 | -1.51 | 0.011 | -1.96 |
| Phylogenetic repulsion | Phylogenetically related species segregate among sites | 0.031 | 5.16 | 0.013 | 2.07 |
| Functional attraction | Functionally similar species aggregate within forest stands | 0.269 | 299.98 | 0.372 | 633.49 |
| Phylogenetic attraction | Phylogenetically related species aggregate within forest stands | 0.265 | 160.35 | 0.252 | 169.54 |

**Table S4.** Summary information for quadrat-scale models quantifying the strength of functional repulsion, phylogenetic repulsion, functional attraction, and phylogenetic attraction in local species co-occurrence patterns (see **PGLMM-3** in Methods). We fit separate PGLMMs for all 257 forest sites and compared models with and without covariance matrices specifying site-level functional repulsion, phylogenetic repulsion, functional attraction, and phylogenetic attraction. The table includes mean standard deviation quantifying the amount of residual variation attributed to functional and phylogenetic repulsion or attraction as well as the number of sites (percent of total sites in parentheses) for which each covariance matrix substantially improved model performance (ΔWAIC greater than 5). See Fig. 5 and Figs. S9-S11.

| Model term | Biological interpretation | Including traits |  | Excluding traits |  |
| --- | --- | --- | --- | --- | --- |
| | | Mean $\sigma$ | No. sites with $\Delta\text{WAIC} > 5$ (%) | Mean $\sigma$ | No. sites with $\Delta\text{WAIC} > 5$ (%) |
| Functional repulsion | Functionally similar species segregate among microsites | 0.059 | 14 (5.4%) | 0.057 | 12 (4.7%) |
| Phylogenetic repulsion | Phylogenetically related species segregate among microsites | 0.055 | 14 (5.4%) | 0.053 | 10 (3.9%) |
| Functional attraction | Functionally similar species aggregate within microsites | 0.114 | 54 (21.0%) | 0.108 | 46 (17.9%) |
| Phylogenetic attraction | Phylogenetically related species aggregate within microsites | 0.145 | 55 (21.4%) | 0.142 | 53 (20.6%) |

**Table S5.** Phylogenetic signal in functional traits

| Trait | Pagel's $\lambda$ | P-value | Blomberg's K | P-value |
| --- | --- | --- | --- | --- |
| Leaf height | 0.520 | 0.013 | 0.054 | 0.016 |
| SLA | 0.666 | 0.003 | 0.051 | 0.061 |
| Leaf C:N | 0.824 | <0.001 | 0.094 | 0.002 |
| Seed mass | 0.982 | <0.001 | 0.953 | 0.001 |

**Figure S1.**
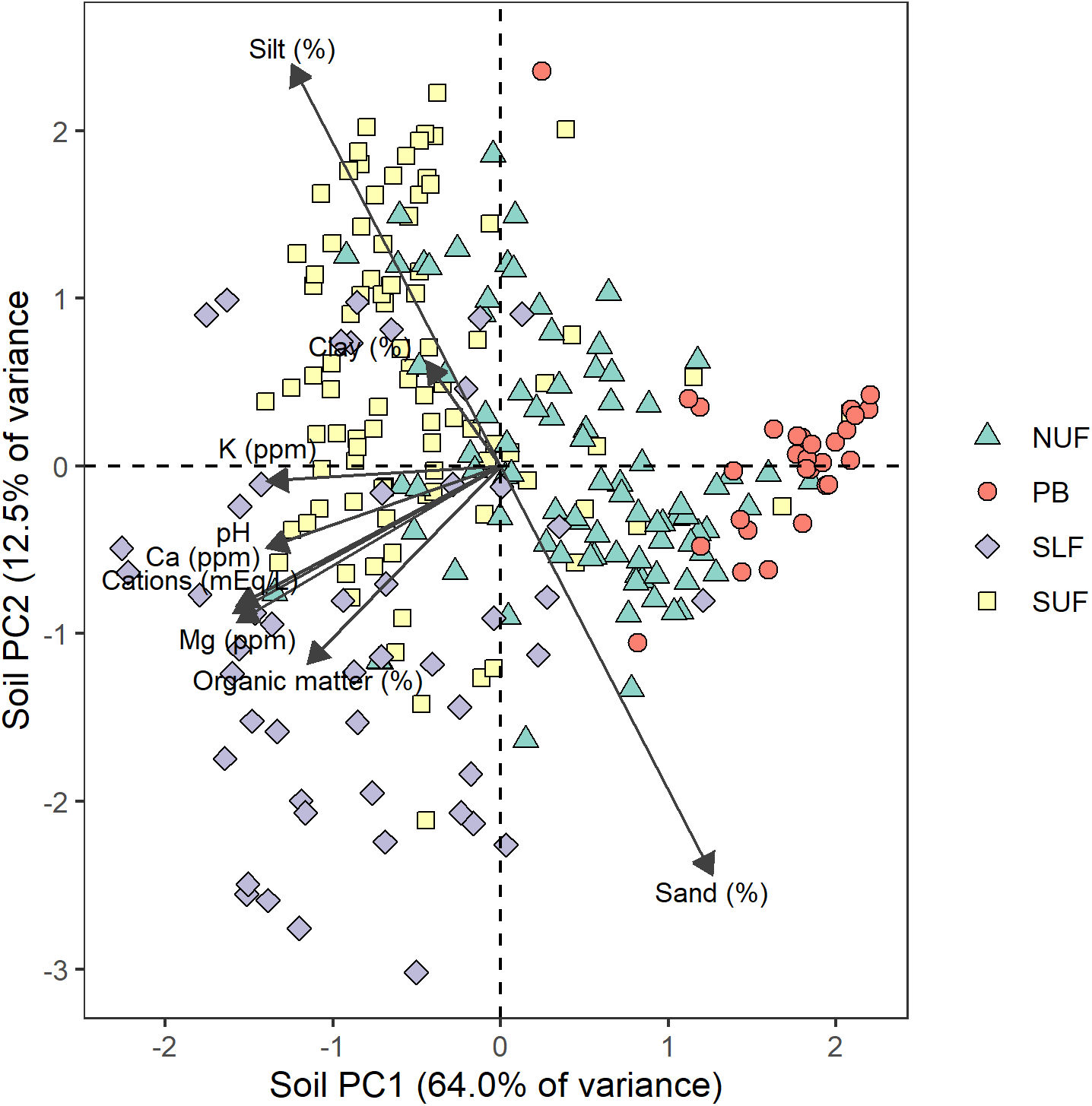
PCA of environmental variables results of principal components analysis illustrating variation in soil characteristics among study sites and community types. We log-transformed soil organic matter content and cation concentrations to improve the linearity of relationships to soil variables. The first principal component (PC1) of the soil PCA accounts for 60.7% of the variance in soil characteristics – primarily the gradient from sandy, nutrient-poor sites to more fertile soils with more silt and clay. The second principal component (PC2) accounts for 13.6% of the variation and is strongly associated with differences in soil organic matter and clay and silt content.

**Figure S2.**
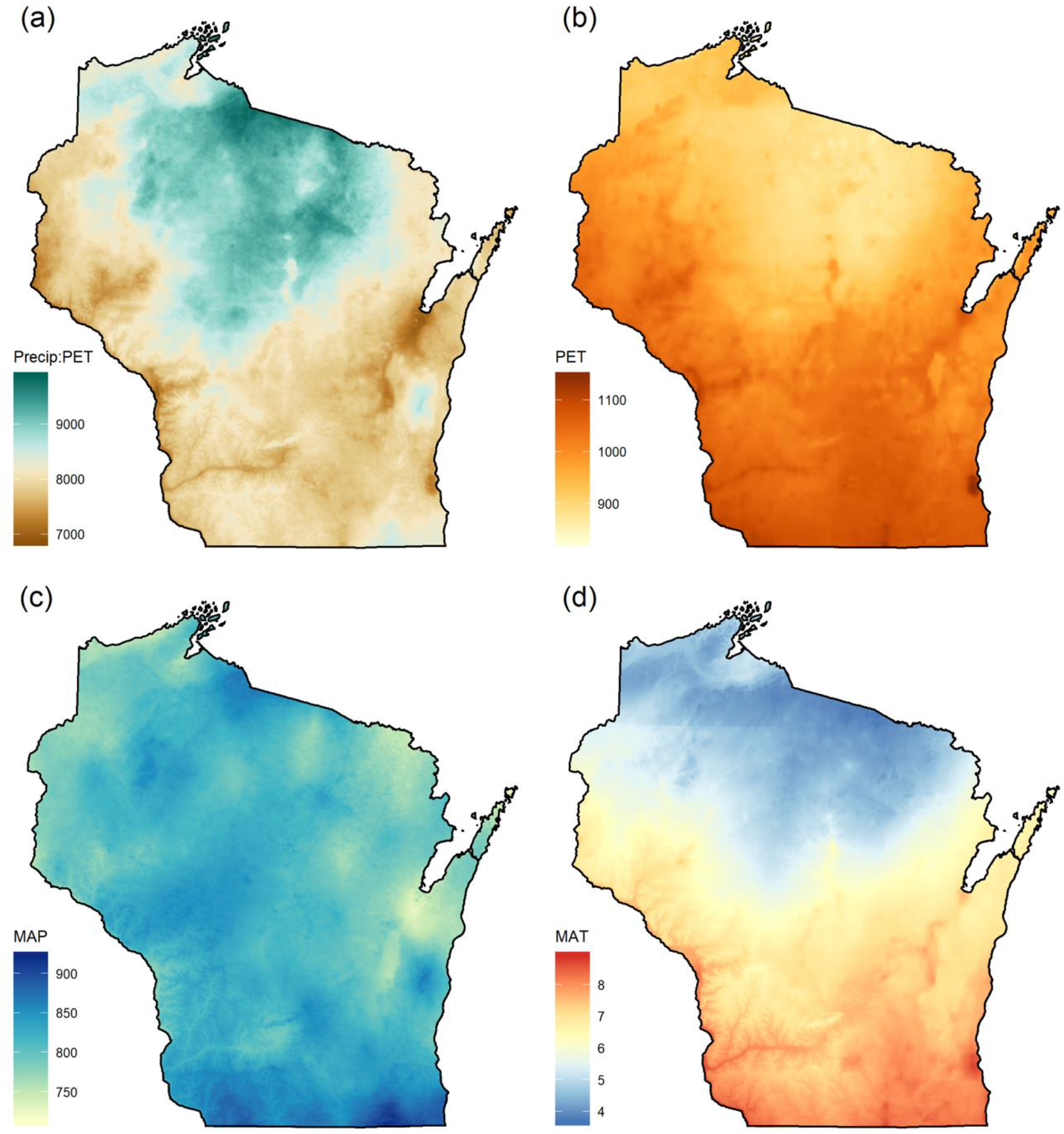
Variation in climate variables across the study area (Wisconsin, USA) including: (a) Precip:PET – ratio of mean annual precipitation to potential evapotranspiration, (b) PET – potential evapotranspiration, (c) MAP – mean annual precipitation (in mm), and (d) MAT – mean annual temperature (°C). These data are derived from 1 km resolution WORLDCLIM climate data from 1970 – 2000.

**Figure S3.**
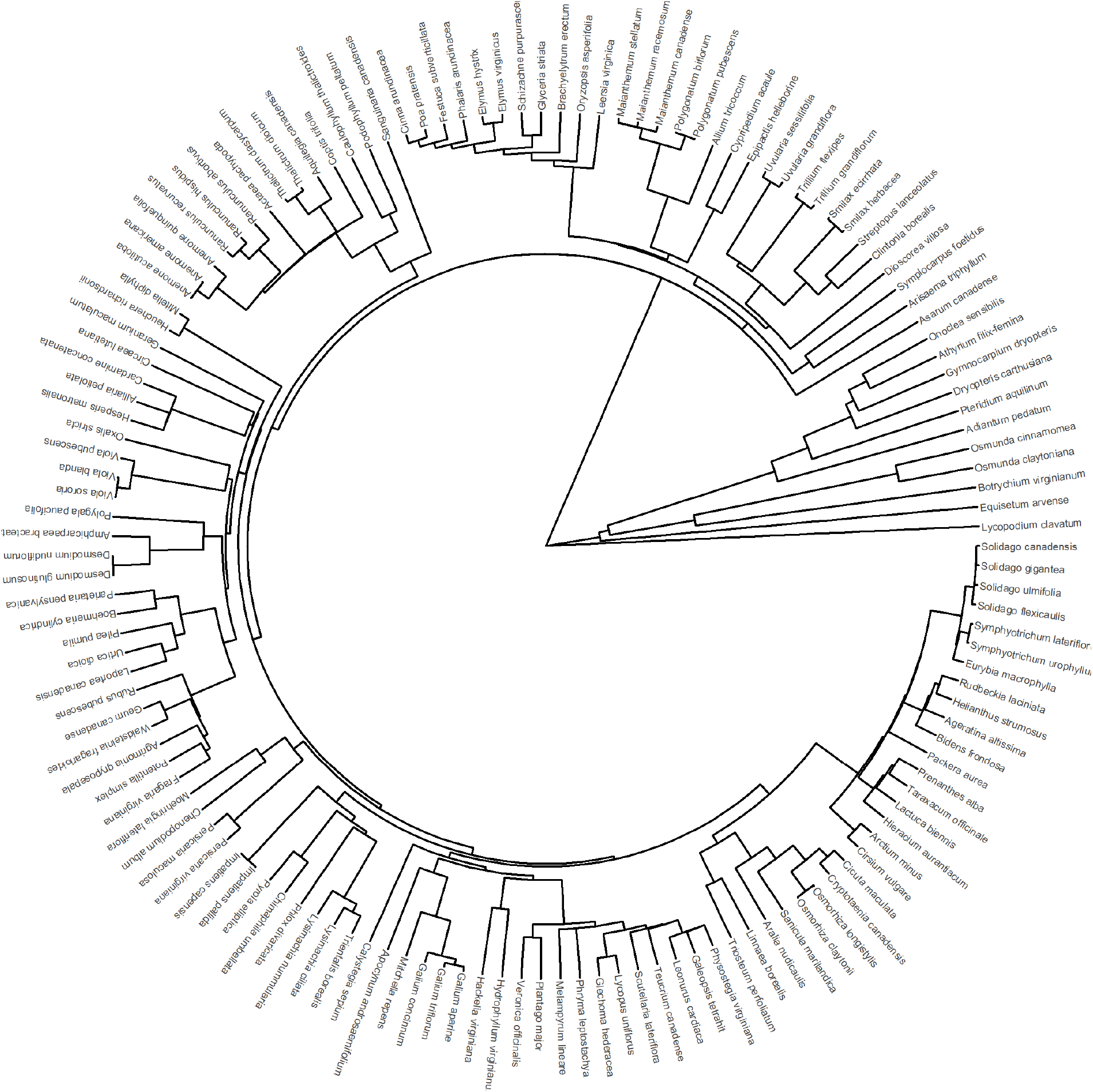
Phylogeny of the 139 herbaceous plant species included in our analysis. See Spalink et al. (2018) for detailed information about the methods used to assemble this tree.

**Figure S4.**
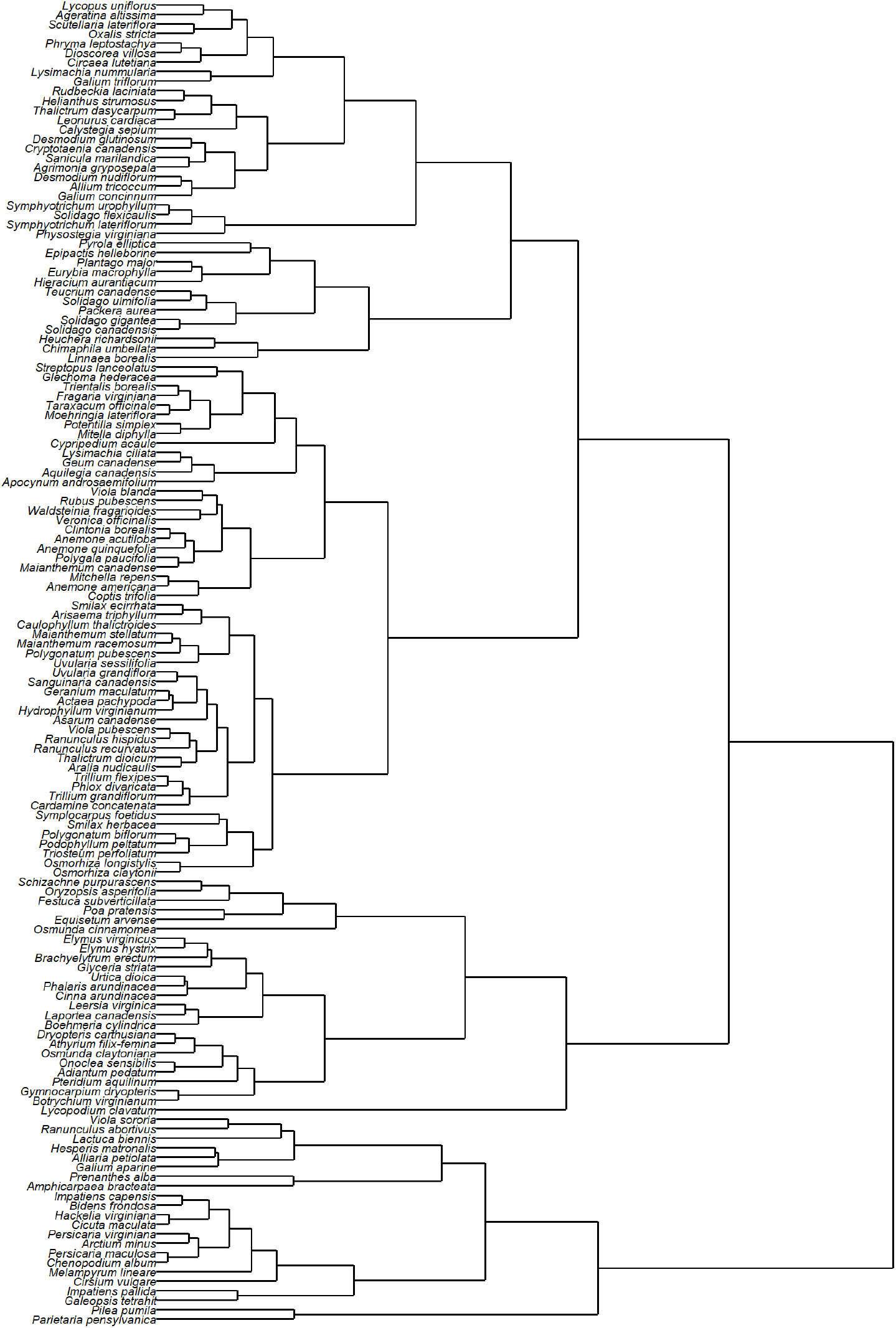
Functional dendrogram illustrating patterns of trait similarity among the 139 focal herb species. The dendrogram was produced using hierarchical clustering algorithm and trait values for leaf height (log-transformed), SLA, leaf C:N, seed mass (log-transformed), species longevity (short-lived versus long-lived), pollination syndrome (biotic versus abiotic), and bloom season (spring, summer, or fall).

**Figure S5.**
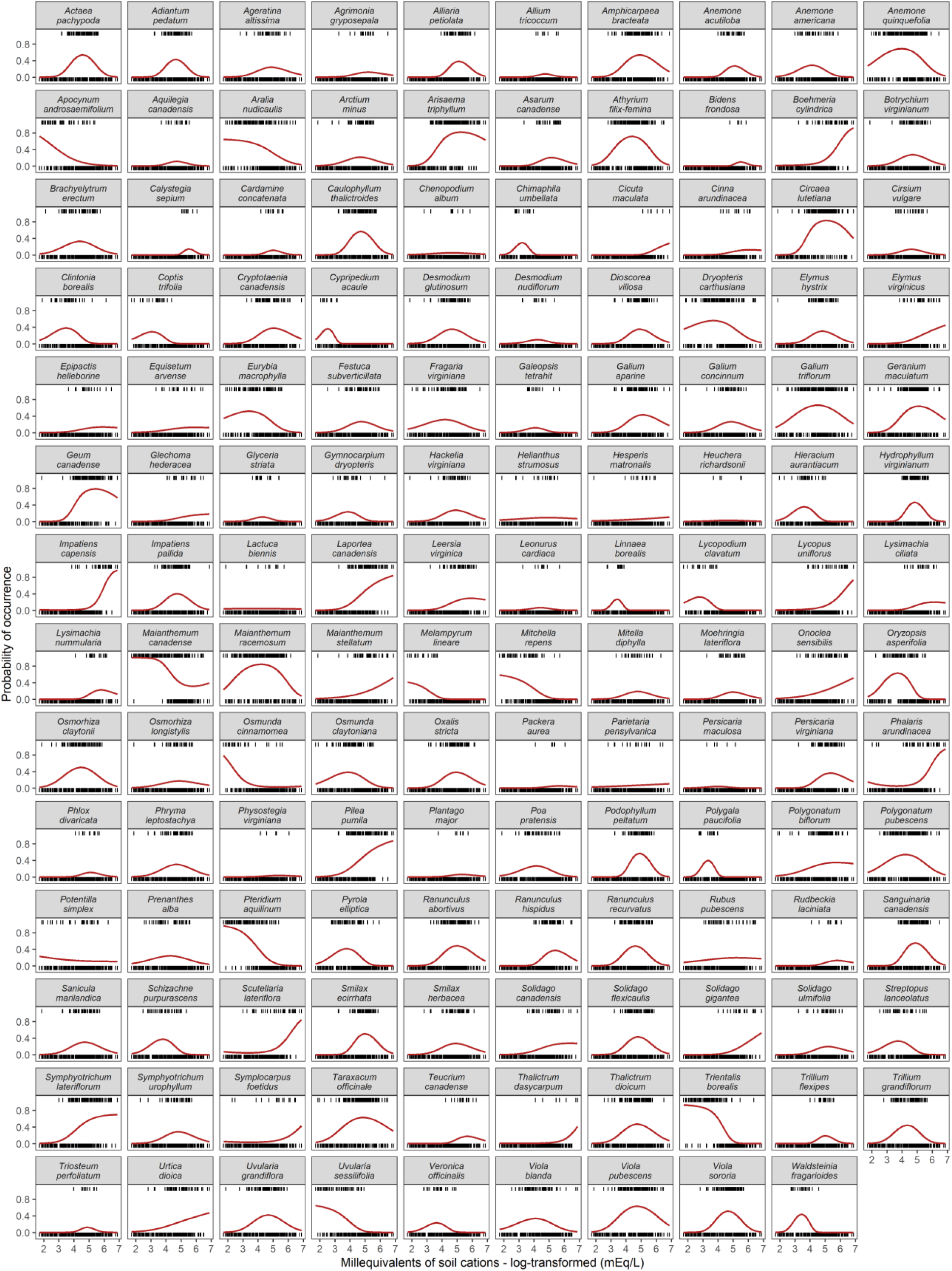
Observed and modeled herb distributions along gradients in soil fertility (log-transformed milliequivalents of leading soil cations: Ca, Mg, and K). Tick marks represent observed occurrences (tick mark at y = 1) and absences (tick mark at y = 0) across the 257 forest stands. Red line illustrates predicted probability of occurrence for each herb species.

**Figure S6.**
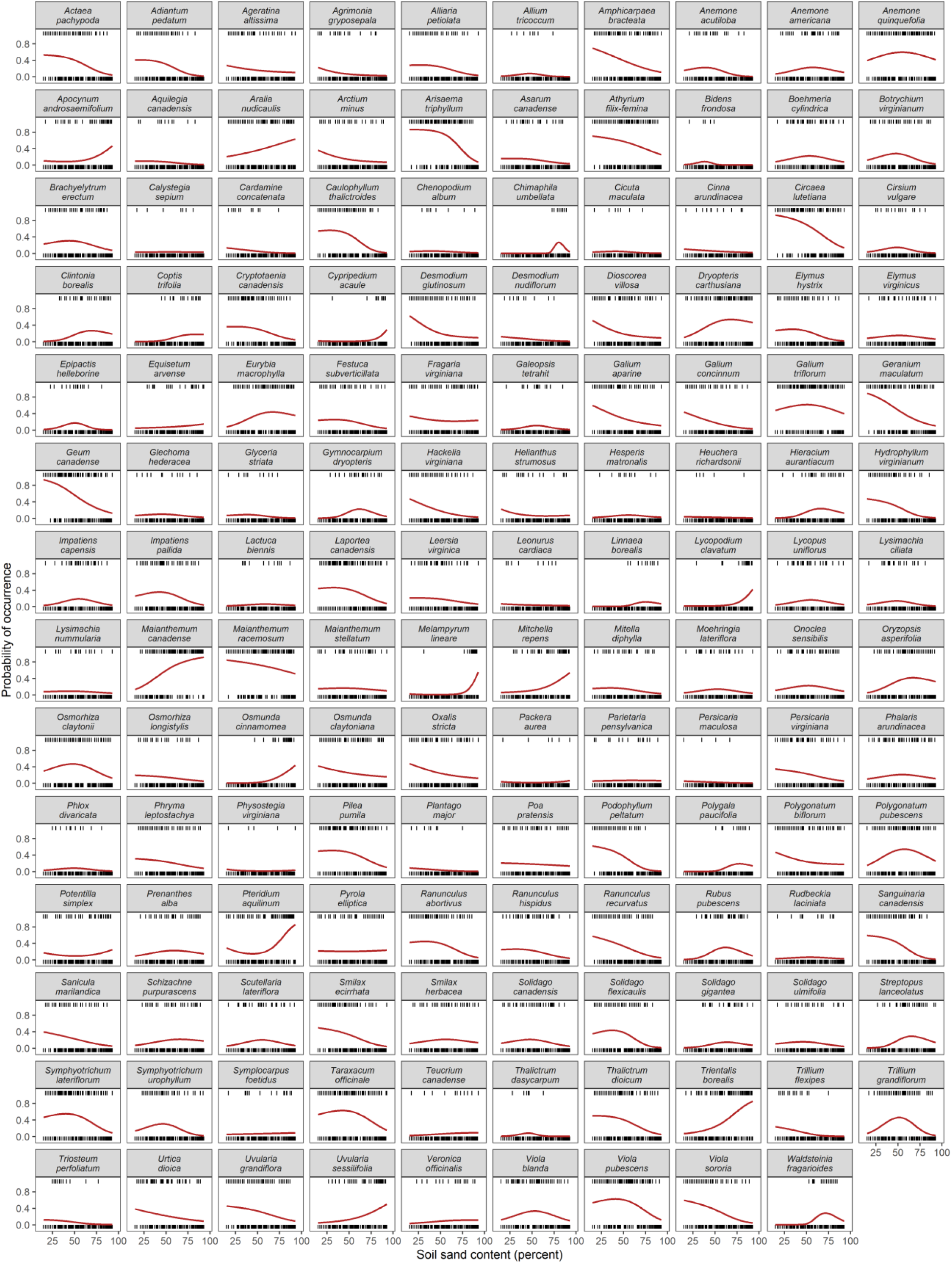
Observed and modeled herb distributions along gradients in soil texture (percent soil sand). Tick marks represent observed occurrences (tick mark at y = 1) and absences (tick mark at y = 0) across the 257 forest stands. Red line illustrates predicted probability of occurrence for each herb species.

**Figure S7.**
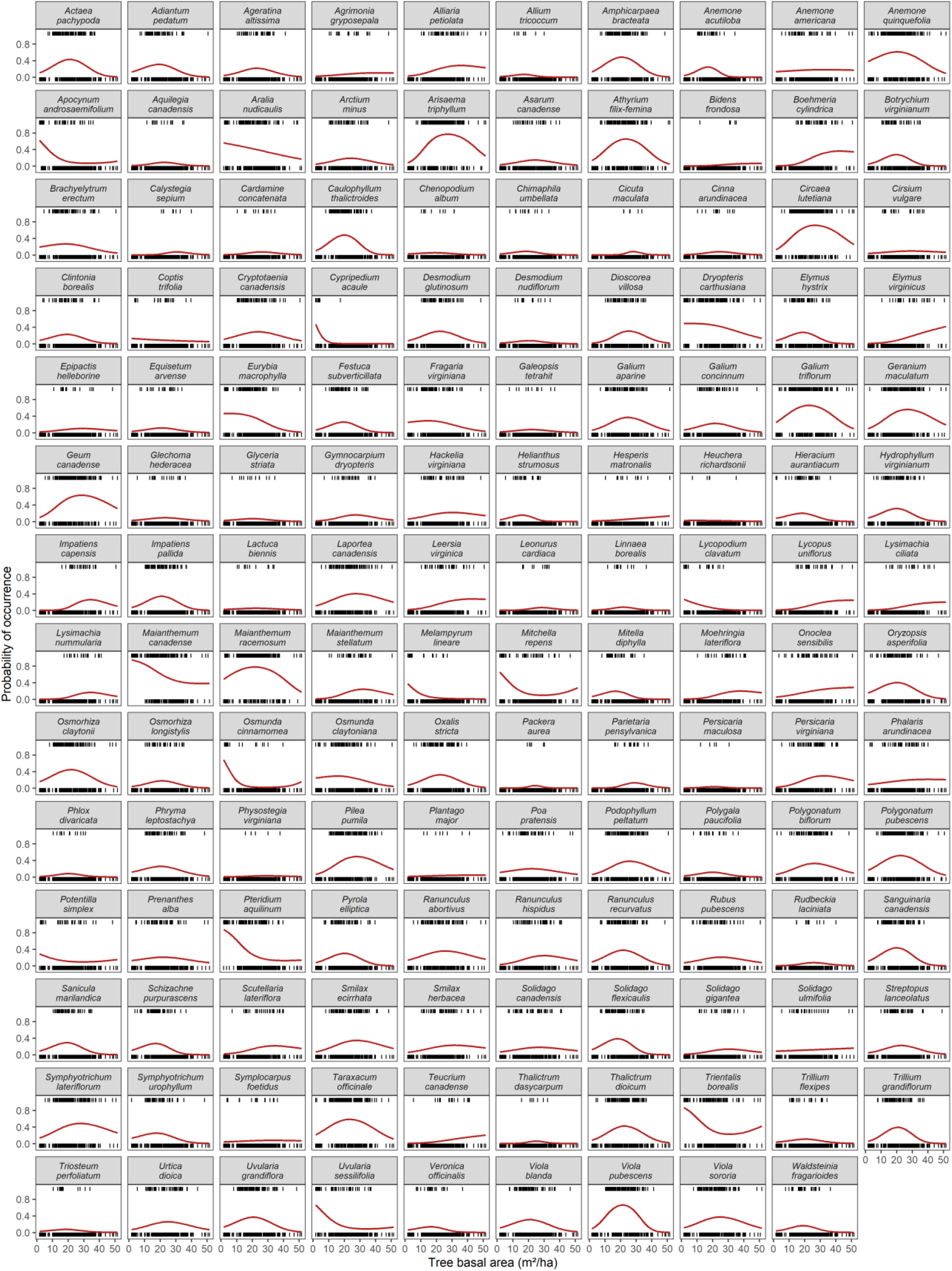
Observed and modeled herb distributions along gradients in tree basal area (m^2^/hectare). Tick marks represent observed occurrences (tick mark at y = 1) and absences (tick mark at y = 0) across the 257 forest stands. Red line illustrates predicted probability of occurrence for each herb species.

**Figure S8.**
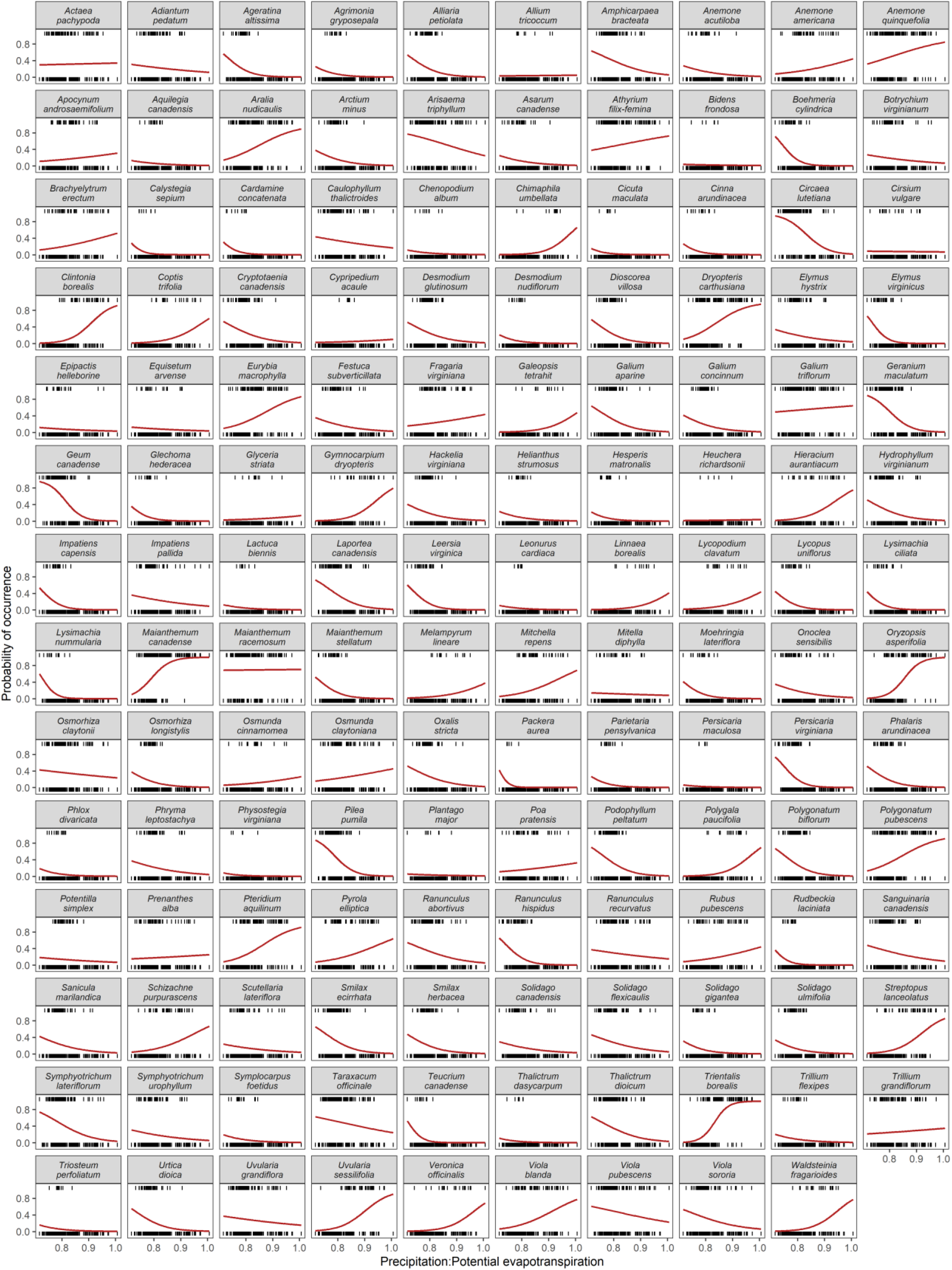
Observed and modeled herb distributions along gradients in Precipitation:Potential evapotranspiration. Tick marks represent observed occurrences (tick mark at y = 1) and absences (tick mark at y = 0) across the 257 forest stands. Red line illustrates predicted probability of occurrence for each herb species.

**Figure S9.**
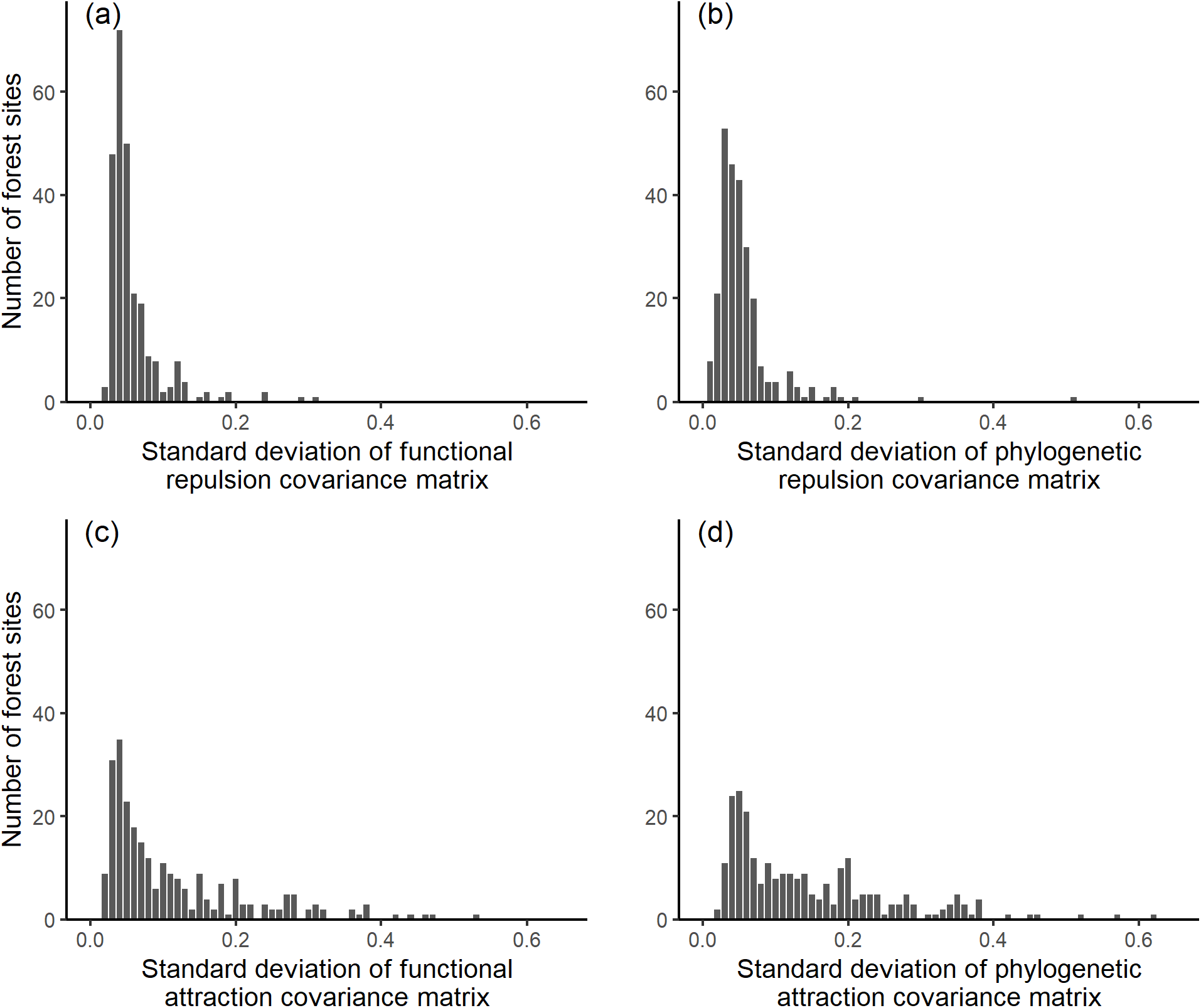
Standard deviation estimates of (a) functional repulsion, (b) phylogenetic repulsion, (c) functional attraction, and (d) phylogenetic attraction for quadrat-scale PGLMMs including trait predictors. We fit separate PGLMMs for all 257 forest stands (see PGLMM-3 in Methods).

**Figure S10.**
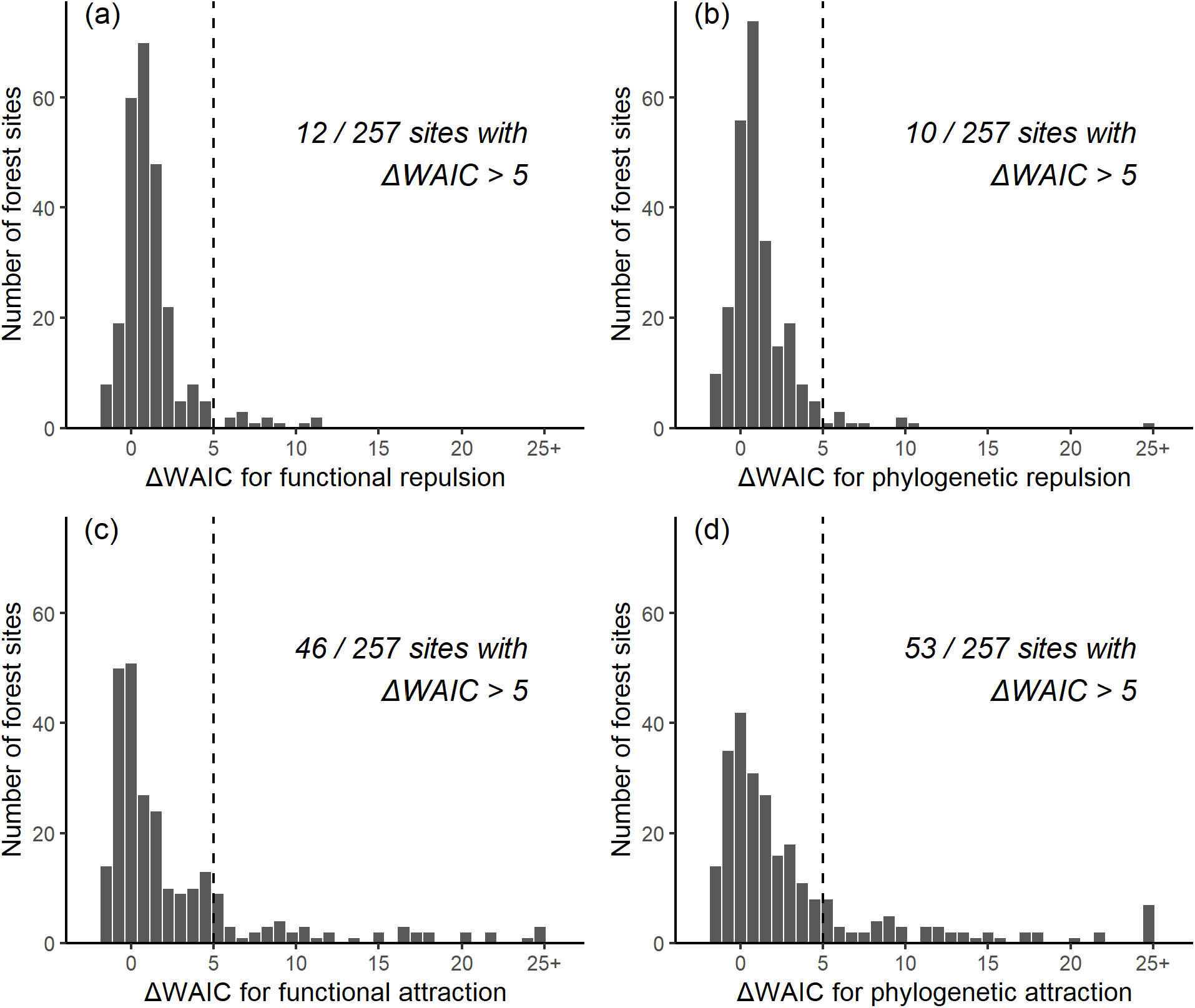
Patterns of (a) functional repulsion, (b) phylogenetic repulsion, (c) functional attraction, and (d) phylogenetic attraction for quadrat-scale PGLMMs excluding trait predictors. These histograms display ΔWAIC values (the differences in WAIC values between PGLMMs with and without covariance matrices specifying repulsion or attraction, see PGLMM-3 in Methods) for all 257 forest sites. Only ΔWAIC values >5 (vertical dashed line) provide evidence of functional or phylogenetic repulsion (spatial segregation among species within sites). Negative ΔWAIC values reflect cases where the penalty for additional complexity is greater than the amount of additional information gained by accounting for functional or phylogenetic repulsion.

**Figure S11.**
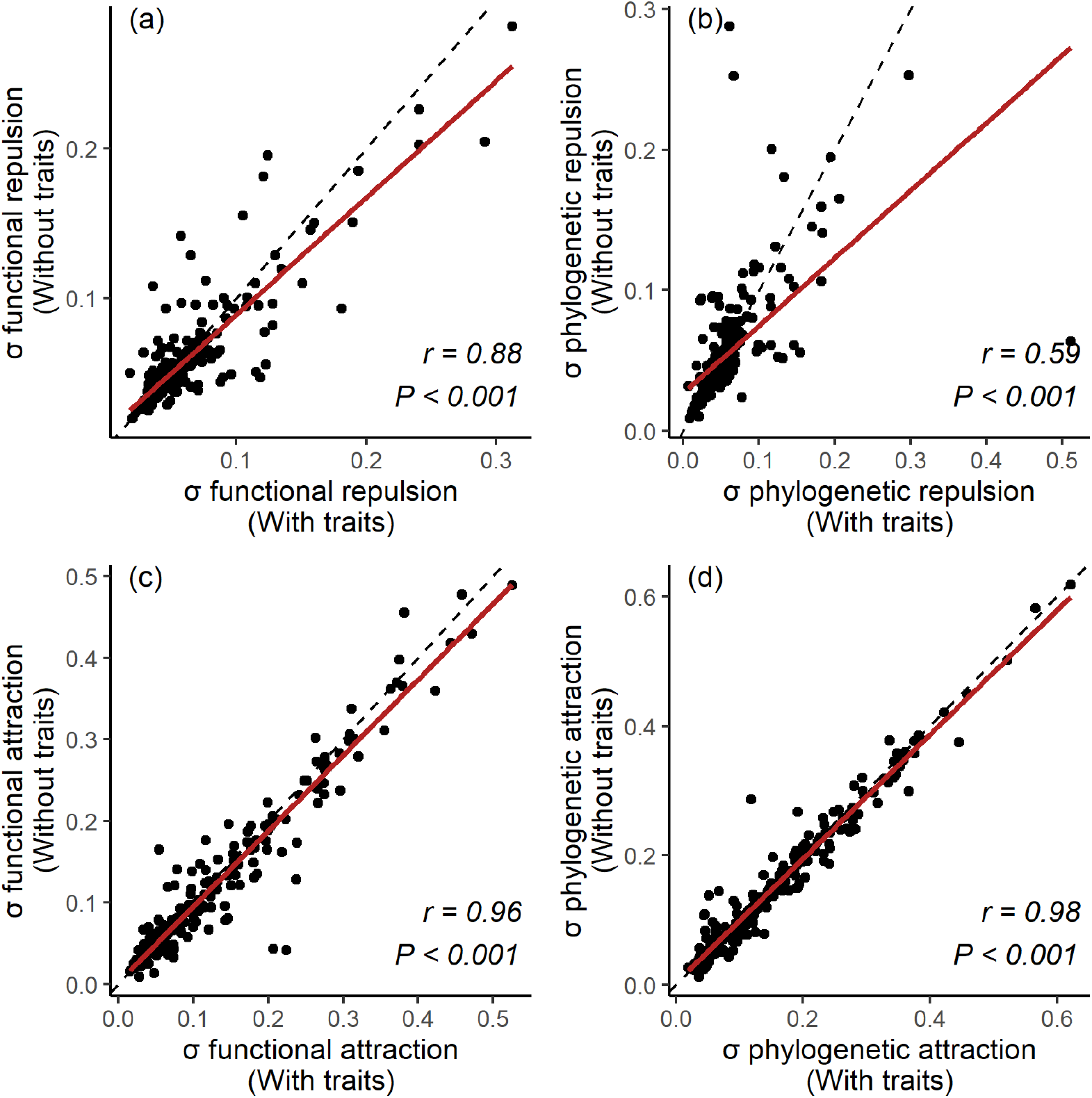
Scatterplots illustrating observed correlation between standard deviation estimates of (a) functional repulsion, (b) phylogenetic repulsion, (c) functional attraction, and (d) phylogenetic attraction for quadrat-scale PGLMMs including and excluding trait predictors. We fit PGLMMs of quadrat-scale species occurrences for all 257 forest sites with trait predictors and without predictors (see PGLMM-3 in Methods). The dashed line illustrates the 1:1 relationship expected if PGLMMs including and excluding trait predictors yield identical estimates of residual standard deviation. The red line illustrates the observed relationship between standard deviation estimates for PGLMMs including and excluding trait predictors. Correlation coefficients and P-values are reported for each scatterplot in the lower right corner.

